# Txn-Txnrd1 system supports redox rewiring during polyaneuploid transition and protects giant cancer cells at new redox homeostasis

**DOI:** 10.64898/2026.08.27.746985

**Authors:** Kinga Kołacz-Milewska, Karolina Gronkowska, Sylwia Michlewska, Markus Absenger, Eleonore Fröhlich, Agnieszka Robaszkiewicz

## Abstract

Polyaneuploid giant cells (PGCC), which occur more frequently in *TP53*-mutant tumors, are recognized as a driver of tumor recurrence and therapy resistance, but the mechanisms supporting their survival remain largely unknown. Our results indicate that polyaneuploid transition and subsequent PGCC maturation in drug-resistant phenotypes are associated with redox rewiring that shifts cellular homeostasis into mild pro-oxidative condition. These are accompanied by increased transcription of genes involved in protection against elevated reactive oxygen species and glutathione-dependent xenobiotic detoxification such as *TXN*, *PRDX2/5*, *GPX1*, and *GSTP1/GSTO1*. Functional studies provided evidence on the crucial role of Txn-Txnrd1 system in maintaining PGCC viability and their adaptation to increased level of reactive oxygen species. Pharmacological targeting of Txn or Txnrd1 as well as their silencing caused a decline in thiol content followed by further redox imbalance, which led to massive death of PGCC. Analysis of clinical datasets revealed direct and relatively strong link between transcription of *TP53* and *TXN* or *TXNRD1*. Tumors with *TP53*^low^/*TXN*^high^ or *TP53*^low^/*TXNRD1*^high^ were associated with considerably poorer patient outcome, whereas elevated transcription of both *TXN* and *TXNRD1* predicted reduced response to chemotherapy in glioblastoma and intestinal cancer. Concluding, Txn-Txnrd1 system enables PGCCs to tolerate pro-oxidative condition, thereby creating a therapeutically exploitable redox vulnerability of these cells, where Txnrd1 emerges as a potential target candidate to overcome PGCC-driven chemoresistance.

**Highlights:**

- Polyaneuploid transition is associated with redox rewiring into more oxidative condition
- Txn-Txnrd1 system is crucial for new redox homeostasis
- Survival of mature PGCC is critically dependent on the Txn-Txnrd1 axis
- *TP53*-low tumors with high expression of *TXN* or *TXNRD1* show poorer clinical outcome

## Introduction

Despite the development of advanced therapeutic strategies such as targeted therapy, immunotherapy and antibody–drug conjugates (1), standard chemotherapy remains the primary treatment option (2,3) or component of combination therapy for many cancer types, which lack effective molecularly targeted therapies. These include, for example, triple-negative breast, non-small cell lung and high-grade serous ovarian cancer, osteosarcoma and numerous other (4). However, diverse adaptive mechanisms that collectively reduce the cytotoxic efficacy of anticancer agents arise in response to chemotherapy, thereby leading to multidrug resistance. This occurs through increased drug efflux mediated by ATP-binding cassette (ABC) transporters, glutathione S-transferase-dependent detoxification, enhanced drug metabolism, more efficient DNA repair, increased tolerance to genomic instability and the recently recognized polyaneuploid transition. The latter mechanism triggered by chemotherapy, radiotherapy, and hypoxic stress (5) gives rise to polyaneuploid giant cancer cells (PGCCs), which are characterized by repeatedly replicated DNA in a form of one enlarged or multiple nuclei (6), and at least a threefold increase in cell size (7). Initially, PGCC were regarded as terminally arrested, biologically inactive cells and beneficial outcome of anticancer therapy (8). However, growing evidence indicates that PGCCs can survive prolonged exposure to therapeutic and environmental stress, enter a dormant state, and subsequently generate proliferative daughter cells through asymmetric division - neosis (8–12). Moreover, PGCCs contribute to intratumoral heterogeneity (5) and are responsible for tumour recurrence (13).

Most studies on PGCC have focused on the mechanisms governing polyaneuploid transition such as endoreplication, which encompasses the endocycle and endomitosis, as well as mitotic slippage, incomplete cytokinesis, cell fusion and cellular cannibalism (5,14). However, specific markers and a comprehensive understanding of epigenomic, transcriptomic, proteomic, and metabolomic changes accompanying polyaneuploid transition, PGCC formation, resilience and depolyploidization remain missing. Very recently polyploidization in breast cancer cell lines was referred as transient therapy-induced senescence (TIS), but dozens of known drug resistance mechanisms offered no explanation for this unique drug resistance pattern (15). Other resistance options include genome redundancy buffering against genetic damage and metabolic reprogramming (16). Cisplatin-surviving polyploid cells derived from prostate cancer cell line PC3 increased their uptake of glucose and glutamine to fuel pentose phosphate pathway, which generates 96% of intracellular NADPH for *inter alia* antioxidant defence, while reducing oxidative phosphorylation (17). On the contrary, polyploid prostate and mammary epithelial cells were reported to be enriched in mitochondria, which also served as the major source of reactive oxygen species (ROS) (18). Despite differences in the proposed mechanisms responsible for oxidant generation, PGCCs have consistently been characterized by a pro-oxidative redox state, but the molecular mechanisms and key redox determinants that enable PGCC survival under persistent oxidative stress remain largely undefined.

Phenotypic transition, cell differentiation and polarization, metabolic plasticity as well as adaptation to stress is associated with dynamic adjustment of redox homeostasis, where alteration in ROS production is followed by fine-tuning enzymatic and non-enzymatic antioxidant systems (19). The observed redox resetting and acquisition of new, more oxidative balance has been linked to breast cancer cell epithelial to mesenchymal transition (20), cancer resistance to radiation or chemotherapy drug such as cisplatin (21), fluorouracil (22), mitomycin C, doxorubicin or paclitaxel (23). Enhanced ROS production can be counteracted via metabolic reprogramming and NRF2/KEAP1, HIF1/2 or FOXO pathway activation, which controls transcription of ROS-scavenging and detoxifying enzymes, drug transporters, glutathione (GSH) and thioredoxin (Txn) antioxidant system (24). GSH acts as an electron donor for glutathione peroxidases (GPXs), which reduce hydrogen peroxide and organic peroxides (25), but also as substrate for glutathione S-transferases, which conjugate GSH to electrophilic compounds, facilitating their detoxification and subsequent elimination from the cell. The thioredoxin system operates in parallel and comprises thioredoxin (TRX), thioredoxin reductase (TrxR) and NADPH (26). Reduced thioredoxin transfers electrons to oxidised proteins, restoring the thiol groups to their normal state, and is then regenerated by thioredoxin reductase (27). This system also supplies electrons to peroxiredoxins (PRDX), which reduce hydrogen peroxide, organic peroxides and peroxynitrite. The thioredoxin system plays a role in maintaining the activity of ribonucleotide reductase and thus supplies the deoxyribonucleotides necessary for DNA repair (28). Therefore, therapeutic potential of targeting several major redox systems, including the thioredoxin/thioredoxin reductase axis (Txn1 – PX-12; Txnrd1 – auranofin, metoxafin and gadolinium), GSH synthesis (GCL – BSO), GSTP1 (canfosfamide/TLK286), and SOD1 (ATN-224) entered clinical trials, while other possible redox vulnerabilities are being intensively explored at the preclinical stage.

Polyaneuploid transition is intrinsically linked to aberration in mitotic division, where repeated rounds of genome replication occur in the absence of cytokinesis. This requires dysregulation of cell cycle-related proteins by their abnormal expression or sub-cellular localization (29). Zhou et al focused on CDC25C as a key molecular factor integrating cell cycle progression with the DNA damage response through Bora, Aurora, PLK1, ATM–Chek2 signaling pathway and p53-dependent checkpoint control. Consistent with the established role of p53 in maintaining genome integrity, PGCC formation has been proposed to occur more readily in *TP53*-deficient or *TP5*3-mutant tumors, and numerous experimental papers made use of *TP53* mutated cancers to generate PGCC for mechanistic studies of polyaneuploid transition and depolyploidization (30–35).

In agreement with these premises, our mouse xenograft model demonstrated extensive PGCC formation in doxorubicin-resistant *TP53*-mutant triple-negative breast cancer (MDA-MB-231; p.Arg280Lys (R280K)) (36), whereas other drug-resistant tumors with the wild-type *TP53* showed little or no evidence of PGCC. To identify transcriptional signatures associated with PGCC formation we performed single-cell transcriptomic profiling of paired non-resistant and drug-resistant cell lines following doxorubicin-induced polyaneuploid transition. Such an approach allowed us to discriminate early and mature PGCC subpopulations from non-polyploid cells as well as to identify molecular markers of PGCCs and potential therapeutic targets. To validate our findings and assess the broader importance of the disclosed Txn–Txnrd1 axis in PGCC-driven resistance, we extended our analyses to cell lines derived from other cancer types with *TP53*^wt^ and *TP53*^mut^ as well as to clinical datasets of triple-negative breast cancer patients.

## 2. Materials and methods

### 2.1 Materials

The triple-negative breast cancer cell line MDA-MB-231 was purchased from Sigma-Aldrich. Breast cancer cell line MCF7, non-small cell lung cancer cell lines NCI-H441 and A549 were from ATCC. BIOFLOAT™ 96-well U-bottom plates were provided by faCellitate (Mannheim, Germany).

DMEM High Glucose with L-glutamine and sodium pyruvate, fetal bovine serum (FBS), and antibiotics (penicillin and streptomycin) were purchased from Biowest (CytoGen, Zgierz, Poland).

Auranofin (TXNRD1 inhibitor), PX-12 (TXN inhibitor), NBDHEX (GSTP1 inhibitor), Polaprezinc (PRDX5 inhibitor), H_₂_DCFDA and monobromobimane were obtained from MedChemExpress (MCE). Triton X-100, DAPI, and resazurin sodium salt were purchased from Sigma-Aldrich (Poznań, Poland). Doxorubicin was purchased from Biokom (Janki, Poland).

Nunc™ Lab-Tek™ Chamber Slides, High-Capacity cDNA Reverse Transcription Kit (cat. no. 4368814), SuperSignal™ West Pico Chemiluminescent Substrate (cat. no. 34580), TRI Reagent™ (cat. no. AM9738), Lipofectamine™ RNAiMAX (cat. no. 13778100), Opti-MEM™ I Reduced Serum Medium (cat. no. 31985070), PageRuler™ Prestained Protein Ladder (10–180 kDa; cat. no. 26617), Pierce™ Protease Inhibitor Tablets, EDTA-free (PIC; cat. no. A32965), PMSF Protease Inhibitor (cat. no. 36978), RNase A (cat. no. R1253), dithiothreitol (DTT), PowerUp™ SYBR™ Green Master Mix (cat. no. A25741), Fluoromount-G™ Mounting Medium with DAPI (cat. no. 00-4959-52), Texas Red™-X Phalloidin (cat. no. T7471), oligonucleotides for Real-Time PCR (CSF2, RPSA, CNIH4, CKLF, EIF1, GAPDH, HPRT, ACTB, and TBP), Goat anti-Rabbit IgG (H+L) Cross-Adsorbed Secondary Antibody, Alexa Fluor™ 546 (cat. no. A11010), Goat anti-Mouse IgG (H+L) Cross-Adsorbed Secondary Antibody, Alexa Fluor™ 488 (cat. no. A11001), Silencer™ Select siRNA for TXN1 (s1), and Silencer™ Select siRNA for TXNRD1 (s755; cat. no. 4427038), Anti-SOD2 monoclonal antibody (cat. no. MA1-106) were purchased from Thermo Fisher Scientific (Warsaw, Poland).

Anti-GSTP1 mouse monoclonal antibody (clone 3F2C2; sc-66000) and siRNA control (sc-37007) were obtained from Santa Cruz Biotechnology.

FITC Annexin V Apoptosis Detection Kit with PI (cat. no. 640914) was from BioLegend.

Redox Homeostasis and Signaling Antibody Sampler Kit (cat. no. 16815), including GPX1 (C8C4) Rabbit Monoclonal Antibody (cat. no. 3286), GPX4 Antibody (cat. no. 52455), Thioredoxin 1 (C63C6) Rabbit Monoclonal Antibody (cat. no. 2429), Thioredoxin 2 (D1C9L) Rabbit Monoclonal Antibody (cat. no. 14907), TRXR1 (D1T3D) Rabbit Monoclonal Antibody (cat. no. 15140), TXNIP (D5F3E) Rabbit Monoclonal Antibody (cat. no. 14715), Prdx1 (D5G12) Rabbit Monoclonal Antibody (cat. no. 8499), Phospho-Prdx1 (Tyr194) (D1T9C) Rabbit Monoclonal Antibody (cat. no. 14041), and Anti-rabbit IgG, HRP-linked Antibody (cat. no. 7074), as well as Histone H3 (D1H2) Rabbit Monoclonal Antibody (cat. no. 4499) were purchased from Cell Signaling Technology (LabJOT, Warsaw, Poland).

SeekOne® DD Single Cell 3′ Transcriptome-seq Kit (Beijing SeekGene BioSciences Co., Ltd., Beijing, China), which includes reagents for reverse transcription, cDNA amplification, library construction and cleanup.

### 2.2 Cell lines, induction of drug resistance and PGCC transition

#### 2.2.1. Cell culture

The MDA-MB-231 cells were initially cultured in L-15 medium supplemented with 15% fetal bovine serum (FBS) and penicillin/streptomycin (50 U/ml and 50 μg/ml, respectively), without CO_₂_ equilibration. After five passages, the cells were adapted to grow in DMEM medium supplemented with 10% FBS and penicillin/streptomycin (50 U/ml and 50 µg/ml, respectively) in 5% CO_₂_. The MCF7, H441 and A549 cell lines were cultured in DMEM medium supplemented with 10% FBS and P/S antibiotics (50 U/ml and 50 µg/ml, respectively).

#### 2.2.2. Induction of drug resistance

Cells were cultured in DMEM supplemented with 10% fetal bovine serum (FBS), 50 U/mL penicillin, and 50 µg/mL streptomycin in a humidified incubator at 37°C with 5% CO_₂_. For the generation of drug-resistant cell lines, cells were seeded into 25 cm² culture flasks and maintained separately for each chemotherapeutic agent. Approximately every three weeks, depending on cell recovery and proliferation, cultures were exposed to gradually increasing concentrations of doxorubicin (0.05–1 µM) or cisplatin (1–10 µM) for 48 h. Following each treatment cycle, cells were washed with PBS and cultured in fresh drug-free medium until they regained normal proliferative capacity before the subsequent round of drug exposure.

#### 2.2.3. PGCC transition

To induce the formation of polyaneuploid giant cancer cells (PGCCs), the cells were seeded onto culture dishes and allowed to adhere for 24 hours. The culture medium was then replaced with fresh medium containing 0.1 μM doxorubicin. The cells were then exposed to this treatment for 72 hours under standard culture conditions (37 °C, 5% CO_₂_). After the drug exposure period, the medium was removed, the cells were washed once with PBS and fresh drug-free medium was added. During doxorubicin treatment, we observed the emergence of PGCCs in the culture, and after 72–120 hours, they constituted an increasing proportion of all cells.

### 2.3 Mouse xenograft model

#### 2.3.1. Preparation of Formalin-fixed Paraffin-embedded tissue

Doxorubicin-resistant MDA-MB-231 cells and cisplatin-resistant A549 cells suspended in Geltrex™ LDEV-Free Reduced Growth Factor Basement Membrane Matrix were injected subcutaneously into 6-week-old female athymic, immunodeficient mice (Crl:NU(NCr)-Foxn1nu; Animalab, Poland). 6 weeks after inoculation the arose tumors were isolated, transferred and stored in PBS with 10% Paraformaldehyde. The entire experiment with animals was performed in the Animal Facility of Faculty of Biochemistry, Biophysics and Biotechnology, Jagiellonian University, Poland, under permission nr 72/2023 issued on 06.04.2023 by the 2nd Local Ethical Committee in Krakow.

Tumors obtained from mice were dehydrated by placing the tissues sequentially in containers with 50%, 70%, 95% and 100% ethanol for 1 hour (each incubation was performed twice). Histological cassettes containing tissues were then incubated in xylene for 1 hour, after which the tissues were transferred to a container with fresh xylene and incubated for another hour. The dehydrated tumors were then suspended in melted paraffin and paraffin blocks were prepared.

#### 2.3.2. Tissue sectioning

Tissue sections were cut using a Leica microtome, paraffin blocks were placed in the microtome and 7-10 uM thick slices were cut from the tissue. The sections were then transferred to a water bath and collected on glass slides. The slides were then placed at 65 degrees Celsius for 10-20 minutes until the paraffin had melted. The slides containing the tissue sections were then placed in an incubator set to 60 degrees Celsius and left there for two hours.

#### 2.3.3. Histological staining

The first step in preparing tissues for histological staining was deparaffinisation. Slides were washed three times in xylene, twice in 100% ethanol, twice in 95% ethanol, twice in 70% ethanol, twice in 50% ethanol and twice in distilled water for five to ten minutes each. For antigen retrieval, we used 1x trypsin solution in PBS for 5 minutes at 37°C, followed by boiling for 5 minutes in a microwave in 10 mM citric acid with 0.05% Tween. Permeabilization and blocking were conducted with 1% (twice for 10 minutes) and 5% (30 minutes) fetal bovine serum in PBS with 0.4% Triton X-100. Then, we stained sections with an anti-Txn and anti-Txnrd1 antibodies in 1% FBS in PBST (diluted 1:400) at 4 °C overnight. The next day, we added a secondary antibody diluted 1:400 for two hours. When we did not stain the proteins, the plasma membrane was stained with CellMask™ Deep Red Plasma Membrane Stain and the cell nuclei were stained with SlowFade™ Glass Soft-Set Antifade Mountant (with DAPI).

### 2.4 Quantification of PGCC

The formation of PGCCs was assessed by flow cytometry and by confocal microscopy.

#### 2.4.1. Flow cytometry

Cancer cells were seeded into a 12-well plate at a density of 100 000 cells per well. After 72h incubation with doxorubicin at a concentration 0.1 µM cells were washed twice with PBS, then fixed in 1% formaldehyde solution in PBS and incubated for 15 minutes at room temperature. After centrifugation, the fixed cells were incubated for 15 minutes (room temperature) in 0.05% Triton X-100 solution. The next step was to centrifuge the cells once more and suspend the pellet in PBS with RNAase A. After a 3-hour incubation, DAPI was added to the prepared cell suspension and flow cytometry measurements were performed using a BD FACSymphony™ A1 Cell Analyzer. Cell cycle analysis was performed with the FlowJo Software tool using FSC-A and SSC-A parameters, and the selected population of polyploid cells was then visualised and clustered on a histogram. Measurement of DAPI fluorescence intensity and shifts on the histogram indicated the DNA content of the cell.

#### 2.4.2. Confocal Microscopy

For confocal imaging cells were seeded on Nunc™ Lab-Tek™ Chamber Slide, at density 10 000 cells per well and treated with doxorubicin. Cells were fixed with 4% formaldehyde solution in PBS at room temperature for 15 min, washed 3x with PBS, incubated for 15 min with 1x CellMask™ Plasma Membrane Stains and again washed at least 3 times with PBS. After washing slides were mounted with SlowFade™ Glass Soft-set Antifade Mountant (with DAPI). The confocal laser scanning microscopy platform TCS SP8 (Leica Microsystems, Germany) with objective 63×/1.40 (HC PL APOCS2, Leica Microsystems, Germany) was used for samples visualization. Samples were imaged with the following wavelength values of excitation and emission: 649 and 666 for Cell Mask and 405 and 430 – 480 nm for DAPI. Cells were classified as PGCCs based on their characteristic morphology, which includes markedly enlarged cell size and the presence of one or more polyploid nuclei. Quantification was performed by counting the number of PGCCs in randomly selected microscopic fields and normalising this figure against the total number of cells. The average nucleus area or volume were measured with Leica Application Suite X (LAS X, Leica Microsystems, Germany).

### 2.5 Quantification of gene expression and protein level

#### 2.5.1. Real-time PCR

Total RNA was isolated using TRI Reagent™ according to the manufacturer’s instructions. RNA concentration was determined using a Quantus™ fluorometer. Complementary DNA (cDNA) was synthesized from total RNA using the High-Capacity cDNA Reverse Transcription Kit (Thermo Fisher Scientific) under the following thermal conditions: 25 °C for 10 min, 36 °C for 120 min, 95 °C for 5 min, followed by a hold at 4 °C.

The expression of genes identified as markers of specific cell subpopulations based on transcriptomic analysis was subsequently quantified by quantitative real-time PCR using PowerUp™ SYBR™ Green Master Mix and gene-specific primers:

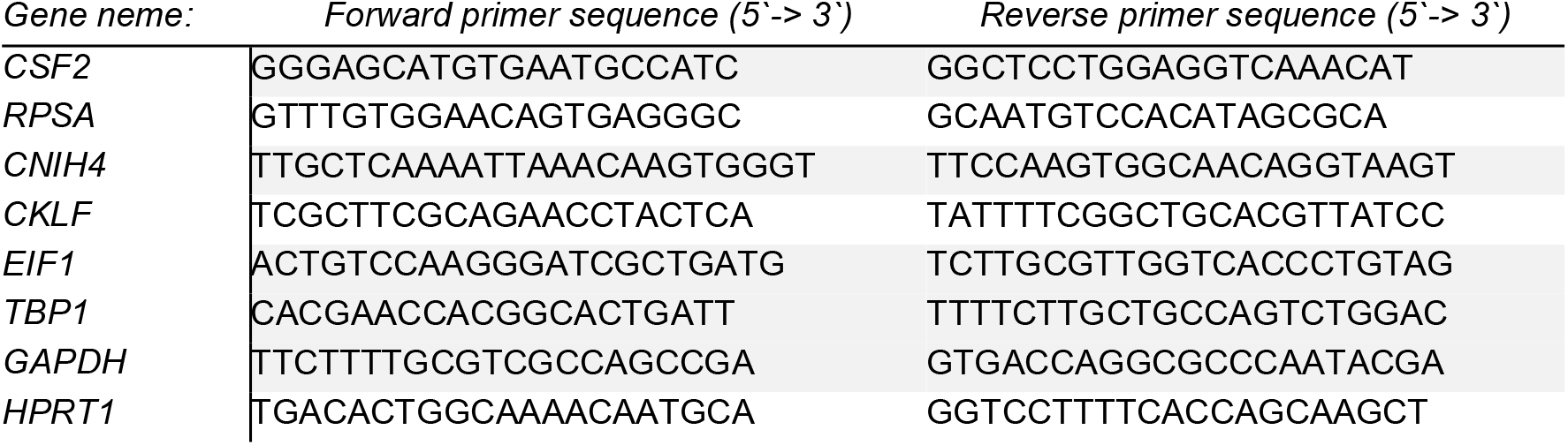

The reaction was run as follow: 50°C, 2 min, polymerase activation: 95°C, 2 min; PCR cycles: denaturation: 95°C,15s; annealing: 57°C, 5s, Elongation: 72°C, 45s). HPRT1, TBP1 and GAPDH (HSKG) were used for normalization and the ratio between the studied gene and HSKG was assumed as 1 for non-treated cells.

#### 2.5.2. Western blot

Cell lysates were suspended in RIPA buffer supplemented with 1 mM PMSF, 1× protease inhibitor cocktail and 1× phosphatase inhibitor. Equal amounts of protein were separated by SDS-PAGE and transferred onto nitrocellulose membranes. The membranes were incubated overnight at 4 °C with primary antibodies (1:5000), followed by incubation with HRP-conjugated secondary antibodies (anti-rabbit, 1:10000; anti-mouse, 1:5000) for 2h at room temperature. Protein bands were visualized using SuperSignal™ West Pico Chemiluminescent Substrate and imaged with the ChemiDoc-IT2 system (UVP, Meranco, Poznań, Poland). Histone H3 served as the loading control.

### 2.6 Measurement of cell viability and apoptosis

#### 2.6.1. Assessment of cell viability

Cell viability was determined using a resazurin-based metabolic assay. The cells were seeded into black 96-well plates and then subjected to one of two treatment schedules. In the first approach, the cells were first exposed to doxorubicin (0.1 µM) for 72 hours. They were then treated with the thioredoxin reductase inhibitor (iTXNRD1 – 0,3 µM), the thioredoxin inhibitor (iTXN - 3µM), the glutathione S-transferase inhibitor (iGSTP1 – 1µM), or polaprezinc (as PRDX5 inhibitor, iPRDX5 – 20 µM) for an additional 72h or after 24-hours doxorubicin pre-treatment the cells were transfected with siRNAs targeting *TXNRD1* (si*TXNRD1*) or *TXN* (si*TXN*) for 72 hours. Following the indicated treatments, cells were incubated with resazurin at a final concentration of 0.0125 mg/mL for 2–3 h at 37°C in a humidified incubator with 5% CO_₂_, protected from light. Fluorescence intensity was measured using a BioTek Synergy HTX microplate reader (Biokom, Poland) with excitation and emission wavelengths of 528 nm and 590 nm, respectively. Background fluorescence, determined from wells containing culture medium and resazurin without cells, was subtracted from all measurements. Cell viability was expressed as a percentage relative to untreated non-resistant control cells, whose fluorescence signal was defined as 100%.

#### 2.6.2. Detection of apoptosis and necrosis using flow cytometry and confocal microscopy

MDA-MB-231 cells were seeded into 12-well plates at a density of 5 × 10^4^ cells per well. Next day cells were treated with 0.1 µM doxorubicin and after 24 h cells were transfected with siRNAs targeting TXN or TXNRD1 for 72h using Lipofectamine™ RNAiMAX Transfection Reagent (Thermo Fisher Scientific) according to the manufacturer’s protocol. Cells transfected with a non-targeting siRNA served as the negative control.

Following the indicated treatments, cells were washed once with PBS and stained with FITC-conjugated Annexin V and propidium iodide (PI) using the FITC Annexin V Apoptosis Detection Kit with PI (BioLegend) according to the manufacturer’s instructions. Samples were analyzed by flow cytometry. FITC fluorescence was detected using 488 nm excitation and 530/30 nm emission, whereas PI fluorescence was detected using 561 nm excitation and 586/15 nm emission. The percentages of viable, early apoptotic, late apoptotic, and necrotic cells were determined based on Annexin V and PI staining.

For confocal microscopy, the cells were seeded onto Nunc™ Lab-Tek™ chamber slides or a 96-well U-bottom BIOFLOAT™ ultra-low attachment plate for spheroids. The cells were then treated with doxorubicin and inhibitors of TXN and TXNRD1 (for 2D cell culture) or siTXN and siTXNRD1 (for 3D cell culture - spheroids). These treatments were carried out at the concentrations and according to the schedules described above. Following treatment, cells were washed once with PBS and incubated with FITC-conjugated Annexin V and propidium iodide (PI) using the FITC Annexin V Apoptosis Detection Kit with PI (BioLegend) for 15 min at room temperature in the dark. Subsequently, cells were fixed with 4% formaldehyde, mounted using SlowFade™ Glass Soft-set Antifade Mountant with DAPI (Thermo Fisher Scientific), and analyzed using a TCS SP8 confocal microscope - 2D cultures (Leica Microsystems, Germany) equipped with a HC PL APO CS2 63×/1.40 oil immersion objective (Leica Microsystems, Germany) and Leica TCS LSI confocal microscope - 3D cultures. Images were acquired using the following excitation and emission settings: 488 nm and 530/30 nm for FITC, 561 nm and 586/15 nm for PI and 405 and 430 – 480 nm for DAPI.

Spheroids from the doxorubicin-resistant MDA-MB-231 cell line were generated using 96-well U-bottom BIOFLOAT™ ultra-low attachment plate (faCellitate GmbH, Mannheim, Germany). The cells were detached by trypsinisation, counted and seeded at a density of 2 × 10 cells/well into individual wells in complete culture medium. The anti-adhesive BIOFLOAT™ polymer coating on the plate surface prevented the cells from adhering to it and promoted spontaneous intercellular interactions, resulting in the formation of spheroids. The plates were then incubated under standard culture conditions (37 °C, 5% CO_₂_) for 21 days. Spheroid formation was monitored using a microscope, and the BIOFLOAT™ U-bottom plate recommendations were followed, meaning that no centrifugation step was performed. Care was taken when carrying out medium changes and subsequent treatments to avoid disrupting the spheroids.

### 2.7 Single-cell transcriptomics

#### 2.7.1 Library preparation and sequencing

Single-cell 3′ RNA-sequencing libraries were generated using the SeekOne® DD Single Cell 3′ Transcriptome-seq Kit (SeekGene, Beijing, China) according to the manufacturer’s instructions. Briefly, MDA-MB-231 non-resistant and doxo-resistant cells exposed to 0.1 µM doxorubicin for 72 h were trypsinized, suspended and 10 000 cells were loaded into the SeekOne® Digital Droplet System for barcoding and incorporation of unique molecular identifiers (UMI). Synthesis of first-strand cDNA was performed directly in the emulsion, and was subsequently recovered, amplified by PCR, subjected to enzymatic fragmentation and end repair, adapter ligation and indexing PCR. The generated sequencing-ready libraries were sequenced on Illumina NovaSeqX PE150 by Genomed S.A. (Warsaw, Poland).

#### 2.7.2 Data preprocessing

Bioinformatic analysis was performed using usegalaxy.org platform (Galaxy version 26.1.rc1). Briefly, FASTQ reads were demultiplexed (Drop-seq, --clipAdapterType – Hamming distance, --soloUMIdedup – CellRanger 2-4, --soloCBmatchWLtype – 1MM_multi, Gene: count reads matching Gene Transcript) and mapped to hg19 with RNA STAR Solo using comprehensive gene annotation model in GTF from GENECODE Release 19 (GRCh37.p13) for slice junction. Barcodes were initially filtered with DefaultDrops method with Expected Number of Cells set at 10 000. AnnData file (h5ad) was imported from DropletUtils 10x Matrices (mtx). Filtering was conducted as follows: pp.filter_genes and min_counts 3; flag_genes - ‘MT-’, pp.calculate_qc_metrics - pct_counts_mito; filter_any - pct_counts_mito; pp.filter_cells - min_genes=200 and max_genes=5000.

#### 2.7.3 Identification of subpopulations and marker genes

All 4 samples (Non-res NT, Non-res + Doxo, Doxo-res NT, Doxo-res + Doxo) were combined with Manipulate AnnData: Concatenate along observation matrix. Data were normalized, logarithmized and saved raw using Scanpy Normalize (pp.normalized_total), Scanpy Inspect and manipulate (pp.log1p), Manipulate AnnData (save_raw), respectively. Higly variable genes were identified with Scanpy filter pp.highly_variable_genes and flavor=’seurat’ and then filtered and scaled with Scanpy Inspect and manipulate (pp.scale) with default parameters. After computing PCA (pp.pca) and variance ratio (pl.pca_variance_ratio) for 30 components, the neighbourhood graph (pp.neighbours) was generated with the following parameters: n_neighbors=10, n_pcs=16, method=’umap’, metric= ‘euclidean’, and then embedded with UMAP (tl.umap). Cells were clustered into subgroups with tl.louvain and flavor=vtraag, resolution=0.4, random_state=0.

Marker genes for louvain subpopulations were identified using tl.rank_genes_group with groupby= ‘louvain’, use_raw=’yes’, n_genes=200, method=’Wicoxon-RankSum’, corr_method=’Benjamini-Hochberg’, reference=’rest’. S and G2-relevant genes were scored using tl.score_genes_cell_cycle and custom gene lists, then regressed out with Scanpy RegressOut.

#### 2.7.4 Pseudotime and trajectories

Single-cell pseudotime analysis and trajectory inference were performed with Scanpy toolkit. Cell clusters were first visualized using force-directed graph with fa layout with Scanpy RunFDG and PlotEmbed (--basis= draw_graph_fa, color=’louvain’, – projection=’2D’). Trajectory inference was computed with Scanpy PAGA and Scanpy PlotTrajectory (-- use-key=’paga’, --layout=’Reingold-Tilford’, --solid-edges=’connectivities’, --root=0).

The force-directed graph embedding was subsequently used for diffusion pseudotime inference to reconstruct cellular trajectories with the followings tools: Scanpy DiffusionMap (n_comps=15), Scanpy ComputeGraph and Scanpy PlotEmbed (-- basis=‘draw_graph_fa’, --color=’ louvain,dpt_pseudotime’, –projection=’2D’).

### 2.8 Analysis of clinical data

Probability of breast cancer patient survival was generated for GDC Pan-Cancer (PANCAN) dataset by creating Kaplan Meier plots for subgroups, which differ in *TP53* and *TXN* or *TP53* and *TXNRD1*. *TP53*^low^ subgroup was assumed for log2(fpkm-uq+1)<2.9, whereas *TXN*l^high^ >7.5, *TXNRD1*^high^ >5.5. The same criteria were used to assess the impact of TP53 expression or the presence of a stop codon in *TP53* on *TXN* and *TXNRD1* gene transcription with box plot.

Breast cancer data described by Yau et al. (2010) served to determine metastasis-free survival for the selected phenotypes. The selected dataset comprised a training cohort of 199 node-negative, adjuvant treatment naive HRneg (including 154 Tneg) breast cancer cases curated from three public microarray datasets. Specific subgroups were stratified based on log2(fpkm-uq+1) as follows: *TP53*^low^ <-1, *TXN*^high^ >1 and *TXNRD*1^high^ > 1.5.

The relationship between *TXN* and *TXNRD1* expression levels and response to chemotherapy was analysed using the online ROC Plotter platform. Patients with glioblastoma and colorectal cancer were included in the analysis. Depending on the clinical outcome data available for each cancer type, response to chemotherapy was evaluated based on Overall survival at 16 months in glioblastoma and Response based on RECIST criteria in colorectal cancer.

### 2.9 Measurement of intracellular redox balance

Reactive oxygen species production and thiol level was quantified with fluorescent probes: H2DCF-DA and monobromobimane, respectively. In brief, cells suspended in PBS were incubated with 50 µM monobromobimane or 10 µM H2DCF-DA at 37 C for 60 min. Monobromobimane-stained cells were fixed with 1% formaldehyde in PBS, permeabilized with 0.1% TritonX-100 in PBS with 1% BSA, incubated with RNAse-A at 37 C for 1 h and stained with 5 µM propidium iodide for 10 min.

Cell fluorescence was measured using BD FACSymphony A1 flow cytometer and the following parameters: excitation 405 nm and emission 450/50 nm for monobromobimane, excitation 488 nm and emission 530/30 nm for H2DCF-DA, excitation 561 nm and emission 586/15 nm for propidium iodide.

To determine the contribution of thioredoxin (TXN) and thioredoxin reductase (TXNRD1) to intracellular ROS and thiol levels, doxorubicin-treated cells were transfected with either siCTRL, siTXN, or siTXNRD1 24 h after drug, and measurements were performed after an additional 72 h. In experiments involving pharmacological inhibitors, the inhibitors were added 24 h before the measurements.

## 3. Results

### 3.1 Drug-resistant *TP53^mut^*cells give rise to transcriptionally heterogeneous PGCCs

While analysing confocal images of drug-resistant tumor sections we identified the formation of giant polyaneuploid cells (PGCC) in xenografts formed from doxorubicin-resistant MDA-MB-231 cells (characterized in (37)), which possess a mutation in *TP53.* Although successive doses of doxorubicin, administered every 7 days for six consecutive cycles, significantly inhibited tumor growth (Fig. 1A-B), histological sections revealed the presence of cells with either single enlarged nuclei or containing more than two nuclei within a cell (Fig. 1C). This phenomenon was markedly less prominent in *TP53^wild-type^* tumor derived from cispl-resistant A549 cells exposed to chemotherapy with cisplatin (Supplem. Fig. 1A-F). To quantify the frequency of PGCCs within the tumors, we first analysed the distribution of DNA content at the single-cell level by measuring DAPI-positive nuclei volume in 3D confocal images. While tumors treated with PBS contained a relatively large fraction of cells in the G1 and S phases of the cell cycle, cells in doxo-resistant tumors treated again with anthracycline showed a marked shift in DNA content toward G2-phase, which is consistent with cell cycle arrest after completion of DNA replication (Fig. 1D). We classified PGCCs as cells with a DNA content exceeding 1.5 fold that of cells in G2-phase (>1.5 × G2 DNA content). Based on this distribution, we estimated the frequency of PGCCs, which constituted approximately 10% of cells in PBS-treated tumors, and increased to approximately 30% after doxorubicin treatment (Fig. 1E). As expected, also the nuclei volume substantially increased (Fig. 1F).

**Figure 1.**
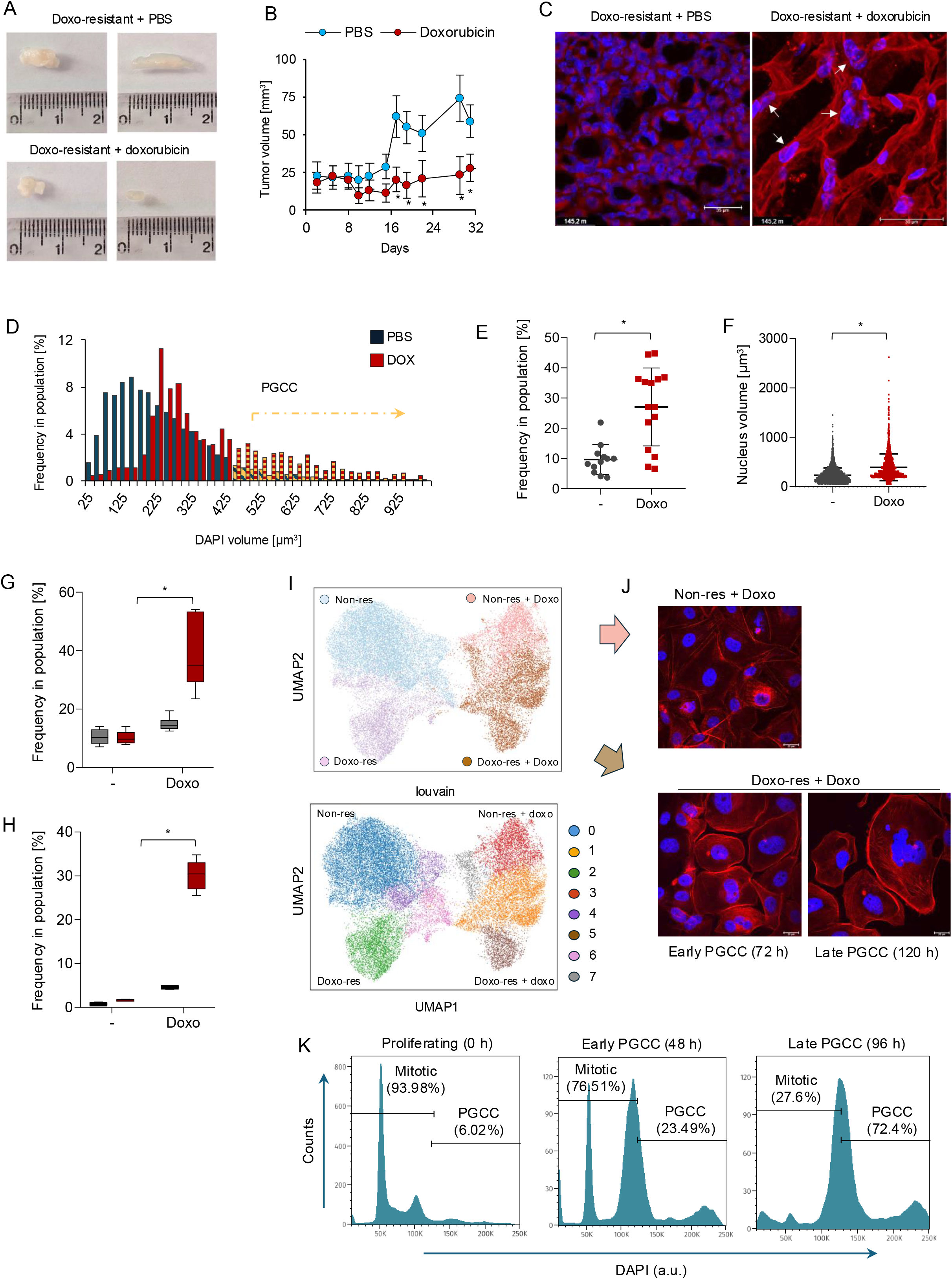
Distinct gene expression-defined subpopulations are present within a PGCC-forming cell line after doxorubicin (A) Representative images of tumors grown and isolated from athymic mice treated with PBS or doxorubicin (5 mg/kg body weight). (B) Tumor growth kinetics (mm³) following PBS or doxorubicin treatment administered every 7 days. (C) Representative fluorescence images of tumor sections from PBS- or doxorubicin-treated mice. Nuclei are stained with DAPI (blue), and actin filaments are stained with Texas Red-X Phalloidin (orange). (D) Distribution of DAPI-stained nuclear volumes determined from confocal microscopy images. Cells with nuclear volume > 3xG1 (highest peak in PBS injected mice) were assumed as PGCC and marked in orange. Quantification of PGCC frequency (E) and nuclear volume (F) in tumor sections. Frequency of PGCC in 2D *in vitro* culture compared between non-resistant and doxo-resistant phenotypes exposed to doxorubicin (0.1 µM) measured by confocal imaging (G) and flow cytometry (H). (I) UMAP projection of integrated single-cell RNA-seq data revealing distinct cellular subpopulations (lower panel) across the four experimental groups (upper panel). (J) Representative confocal images illustrating nuclear enlargement during PGCC formation in 2D culture at the indicated time points. Cells were stained with DAPI (nuclei) and Texas Red-phalloidin (F-actin). Non-resistant cells are shown for comparison of nuclear size. (K) Flow cytometric analysis of DNA content distribution during PGCC formation. Cells with DNA content within the subG1–G2 range were categorized as mitotic, whereas cells with DNA content greater than 2.5x G1 peak in untreated cells were classified as PGCCs.

While considering doxo-resistant MDA-MB-231 cells as a model for testing PGCC development and features, we observed very similar yield of polyploid transition in 2D culture when quantifying their frequency by confocal imaging (Fig. 1G) and flow cytometry (Fig. 1H). These findings suggest that at least in this particular case the mode of cell growth has a relatively minor impact on PGCC formation. To determine whether *TP53* status might be associated with susceptibility to induction of polyaneuploidy, we quantified PGCC formation following doxorubicin treatment in additional drug-resistant cell lines cisplatin-resistant MDA-MB-231 cells, cisplatin-resistant H441 cells, doxorubicin-resistant MCF7 cells (Supplem. Fig 2A) as well as previously characterised cisplatin-resistant A549 cells. The considered anthracycline failed to trigger polyaneuploid transition in chemoresistant *TP53^wt^* genotypes, but substantially elevated frequency of PGCC was found in other *TP53^mut^*phenotypes such as cisplatin-resistant H441 or MDA-MB-231 (Supplem. Fig. 2B).

**Figure 2.**
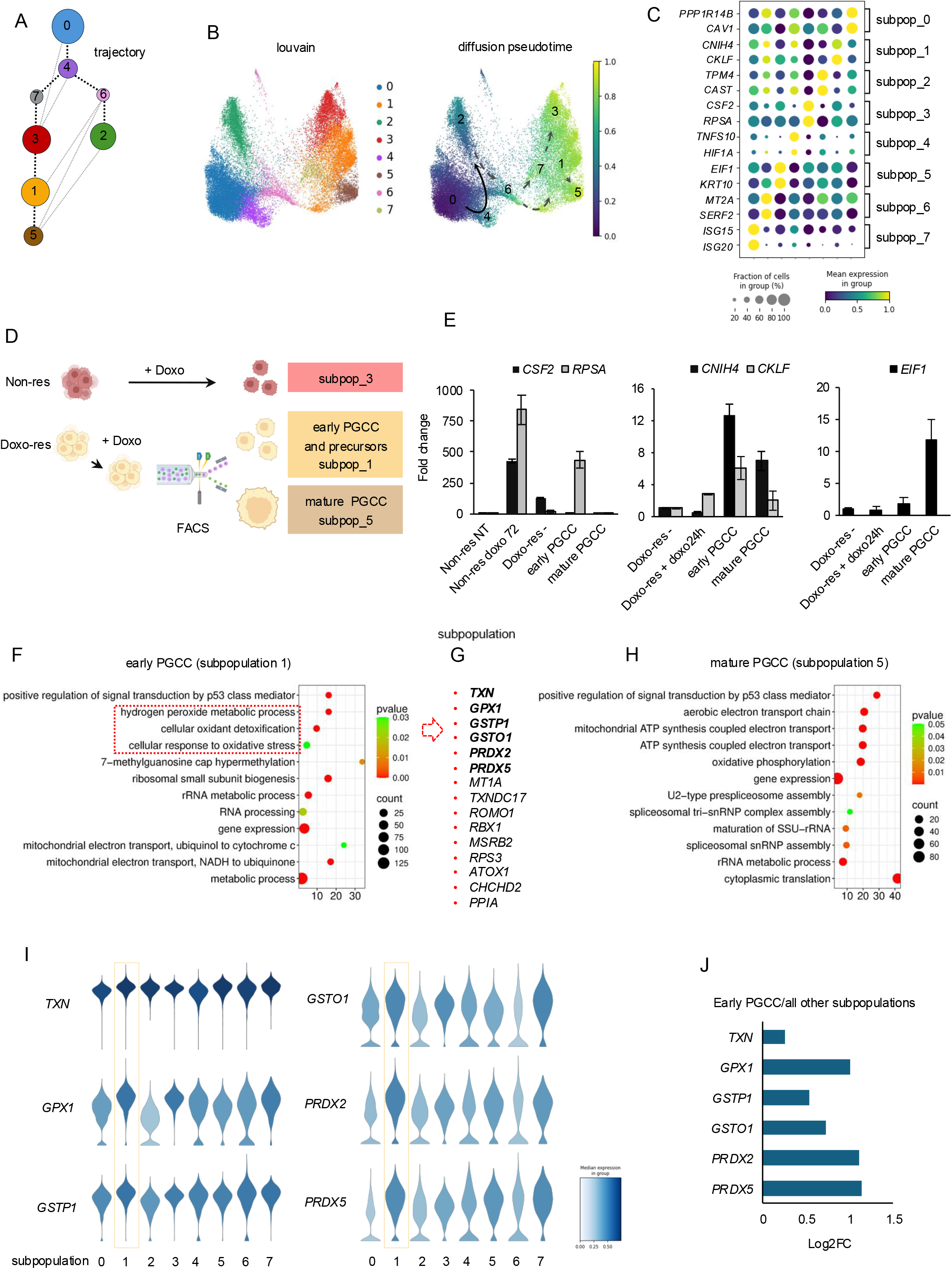
Single-cell transcriptomic analysis revealed the upregulation of genes involved in thioredoxin- and glutathione-dependent detoxification pathways in early PGCC (A) Trajectory inference using PAGA form integrated single-cell transcriptomic data. Cells are colored according to louvain. (B) Diffusion pseudotime trajectory showing the distribution of louvain-defined cell subpopulations. (C) Dot plot showing the expression of representative marker genes across Louvain-defined cell subpopulations. Dot size represents the proportion of expressing cells, and color intensity indicates relative gene expression. (D) Schematic overview of the cell sorting strategy used to separate early PGCCs/PGCC precursors from mature PGCCs according to flow cytometry forward scatter (FSC) and side scatter (SSC) parameters. (E) Real-time PCR validation of selected marker genes for subpopulations 1, 3, and 5 in doxorubicin-treated non-resistant cells as well as in doxo-resistant sorted cells. Gene Ontology (GO; biological process) enrichment analysis of genes differentially expressed in early PGCC/subpopulation 1 (F) and late PGCCs/subpopulation 5 (H). (G) List of genes for “cellular oxidant detoxification” GO term. (I) The six genes highlighted in bold were selected for further comparative analysis of their expression across all identified cell subpopulations using violin plots (I), and against average expression in all other subpopulations (J).

In search for specific markers or targetable factors in PGCC we performed profiling of single-cell transcriptomes of untreated and doxorubicin-treated non-resistant and drug-resistant MDA-MB-231 cells, which indicated that all four samples exhibited distinct gene expression profiles of at least a subset of genes (Fig. 1I, upper panel). Louvain clustering revealed the presence of three clearly distinguishable subpopulations: 1, 5 and 7 in the doxorubicin-treated drug-resistant sample (Fig. 1I, lower panel). Importantly, doxorubicin-induced changes in gene transcription profiles varied in non-resistant and doxo-resistant phenotypes, thereby suggesting their substantially different transcriptomic responses to anthracycline.

To align cellular or nuclear morphology with transcriptomic discrepancies we examined cells by confocal microscopy following the same duration of doxorubicin exposure as used for the single-cell RNA sequencing experiment (Fig. 1J). Under these conditions, doxo-resistant culture was characterized by the mixture of cells with normal and increased nuclear size (early PGCC or PGCC precursors) when compared to culture of non-resistant cells. In extended culture (120 h) giant cells became readily apparent in the resistant population, which is consistent with the formation of mature, late PGCCs. Flow cytometric quantitative analysis of cells with the fluorescently stained DNA revealed a progressive increase in cellular DNA content during prolonged cell growth after doxorubicin treatment (Fig. 1K). Rather than forming a distinct PGCC population, cells exhibited a continuous distribution of DNA content, indicating ongoing genome replication and the generation of cells with probably heterogeneous ploidy, what is consistent with a polyaneuploid rather than a uniformly polyploid phenotype.

Summarizing, in the culture of *TP53^mut^* drug-resistant MDA-MB-231 line, polyploid giant cancer cells emerge as a heterogeneous population characterized by distinct gene expression profiles and the coexistence of both early and mature PGCCs.

Data distribution normality was assessed using the Shapiro–Wilk test. Data in (B), (G) and (H) were analysed using two-way ANOVA. Differences between groups in (E) was analysed using an unpaired t-test, whereas the Mann–Whitney U test was used in (F). Statistically significant differences are indicated by *p* < 0.05 (*).

### 3.2 Early PGCC are characterized by high expression of genes involved in the maintenance of redox homeostasis

To test if transcriptional diversity results from coexistence of early and mature PGCC in drug-resistant *TP53^mut^* cells we reconstructed the cell-state progression based on gene expression profiles in particular subpopulations (Fig. 2A–B). While assuming non-resistant untreated cells (subpopulation 0) as a starting point, trajectory analysis (Fig. 2A) and diffusion pseudotime (Fig. 2B) revealed a branching transcriptional response to doxorubicin rather than a single linear progression. Similarly in both cases, subpopulation 4 emerged as the earliest, major transitional state and the immediate response to doxorubicin (Fig. 2A). Central location of subpopulation 3 in the analysis of trajectory suggested a major intermediate state from which multiple transcriptionally distinct populations emerge, whereas in pseudotime analysis subpopulation 3 consisted of doxorubicin-treated non-resistant cells, represented rather a distinct drug-response state. The apparent inconsistency between the pseudotime progression and the connectivity of subpopulations 1, 5, and 3 may indicate that this connection reflects shared transcriptional responses to doxorubicin rather than a direct developmental relationship. However, the consistency in the progression from subpopulation 1 to 5 prompted us to hypothesise that these two cell states correspond to early and late/mature PGCC.

To test this, we first identified specific markers of louvain-clustered subpopulation (Fig. 2C), and then employed fluorescence-activated cell sorting (FACS) to separate cells according to their size and granularity (Fig. 2D). In accordance with the original assumptions, non-resistant cells treated with doxorubicin expressed higher level of markers typical for subpopulation 3, namely *CSF2* and *RPSA* (Fig. 2E). Doxo-resistant cells, which exhibited markedly higher FSC and SSC parameters, showed elevated expression of *EIF1 -* the subpopulation 5 marker, whereas cells that still contained an approximate single nucleus and size of non-resistant cells were characterized by higher expression of *CNIH4* and *CKLF -* markers of the subpopulation 1. This confirms that at least two transcriptionally distinct subpopulations in doxo-resistant cells exposed to doxorubicin correspond to early and late form of PGCC. This led us to the conclusion that subpopulation 1 corresponds to early PGCC, whereas subpopulation 5 to mature PGCC.

Functional association analysis of highly variable genes in mature PGCCs indicated a metabolic shift toward mitochondrial oxidative phosphorylation, alterations in rRNA metabolism and cytoplasmic translation as well as enhanced signal transduction via p53 pathway (Fig. 2H). Interestingly, in early PGCC a slightly broader spectrum of altered intracellular processes was observed, including enhanced non-coding RNA maturation and stability by 7-methylguanosine cap hypermethylation, as well as activation of antioxidant defence mechanisms, particularly those responsible for hydrogen peroxide detoxification (Fig. 2F). The list of genes linked to these GO terms included phase II detoxification enzymes: glutatione S-transferases, peroxiredoxines 2/5, glutathione peroxidase 1 and tioredoxin (Fig. 2G). Expression of these genes across the identified subpopulations reached the highest values in subpopulation 1 and, for many genes, also in subpopulation 7 (Fig. 2I). In contrast, in subpopulation 5, only thioredoxin (TXN) appeared to be expressed at a level comparable to that observed in subpopulation 1. The magnitude of transcriptional upregulation compared to all other subpopulations revealed the strongest overexpression of peroxiredoxins and glutathione peroxidases in subpopulation 1, whereas thioredoxin exhibited the least pronounced increase in expression (Fig. 2J).

In summary, the activation of multiple genes involved in antioxidant defence is an early event during polyaneuploid transition, whereas mature PGCCs retain only a subset of these adaptations, with sustained thioredoxin expression.

### 3.3 PGCC formation and maturation is associated with shift in redox homeostasis to pro-oxidative condition

To validate the transcriptional upregulation of several genes involved in the redox response and glutathione-dependent detoxification in early PGCCs, we monitored temporal changes in their and other antioxidant protein levels following doxorubicin treatment (Fig. 3A). Gstp1, but also in Txnrd1, Prdx1 and Gpx4 increased within 24 h after drug, whereas substantial increase in Gpx1 and Txn1 levels was detected only in late PGCCs, thereby suggesting a temporal delay between mRNA and protein accumulation. The abundance of proteins, whose transcripts were not upregulated in the single-cell dataset such as Txn2, Txnrd1, Prdx1, Gpx4, indicate that their expression may be regulated post-translationally, allowing a rapid adaptive response to oxidative stress.

**Figure 3.**
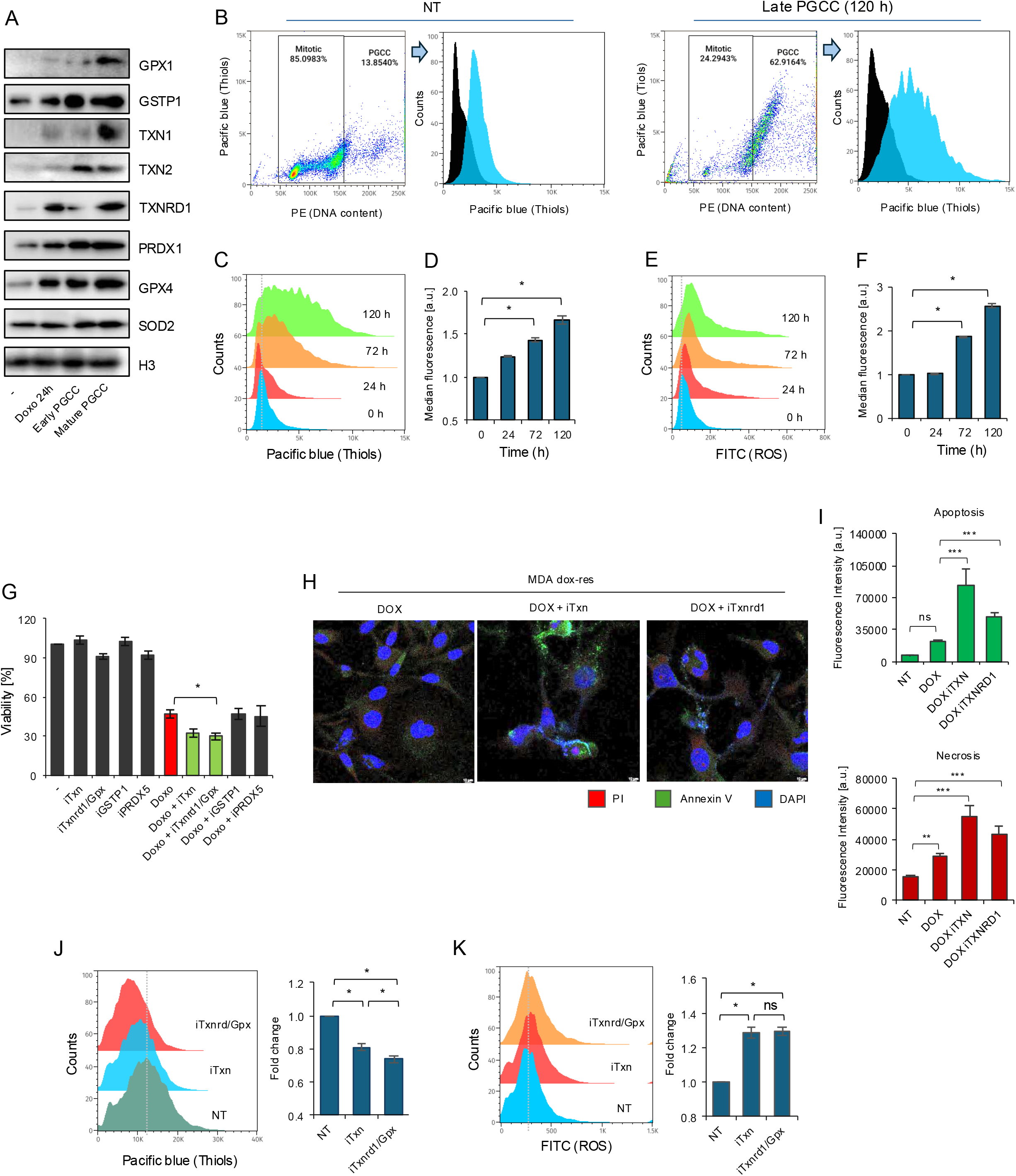
PGCC development shifts redox homeostasis into mild pro-oxidative condition (A) Western blot comparison of selected antioxidant and detoxification proteins in a timecourse of PGCC formation and maturation. (B) Representative analysis of intracellular thiol levels in cells with baseline DNA content and mature PGCCs. Left dotplots indicate DNA (PE) distribution in cell population and allow for discrimination of mitotic cells as well as PGCC, whereas histograms on the right correspond to thiol level (Pacific blue) in both subpopulations. DNA was stained with propidium iodide, whereas thiols with monobromobimane. (C) Histogram representation of the shift in thiol content in the whole cell population at various time points after doxorubicin and (D) quantification, where bars correspond to fluorescence intensity normalized to untreated cells (NT). (E-F) Similar approach was employed to measure distribution of green fluorescence (H2DCF-DA) that contributes to ROS content inside cells. (G) Cell viability was measured with resazurin assay after the following treatments: enzyme inhibitors were added for 72 h to drug untreated cells or to already forming PGCC (72 h after doxorubicin). (H) Representative confocal images of doxorubicin-resistant MDA-MB-231 cells following Txn or Txnrd1/Gpx1 inhibition. iTxn and iTxnrd1 were added 72 h after doxorubicin. DAPI (blue) indicates nuclei, Annexin V–FITC (green) indicates apoptotic cells and propidium iodide (red) indicates necrotic cells. (I) The graphs show the mean fluorescence intensity of FITC (Annexin V; apoptosis) and PI (propidium iodide; necrosis), expressed in arbitrary units (a.u.). Quantification was performed using Leica Application Suite X - LAS X software. (J-K) Thiol and ROS levels were measured as described in (C-F), but iTxn and iTxnrd1/Gpx1 were added 24 h prior to analysis to cells already treated with doxorubicin for 72 h.

Bearing in mind the altered expression profile of antioxidant enzymes, we monitored total thiol content and hydrogen peroxide level in time after induction of polyaneuploid transition. First, to discriminate non-PGCC (entitled mitotic) from PGCC we made use of cell double staining with monobromobimane for thiols and ethidium bromide for DNA content (Fig. 3B). Compared with mitotic cells, PGCCs present in the untreated population already exhibited elevated intracellular level of free thiol groups, which increased even further after doxorubicin treatment. While following cells in time, a gradual increase in intracellular thiol levels was observed, reaching statistical significance 72 hours after doxorubicin treatment, what coincided with the coexistence of both early and mature PGCCs in cell population (Fig. 3C-D). Two non-mutually exclusive explanations or this phenomenon can be considered: a shift in redox balance toward a more reducing state or an increase in total thiol content simply reflecting the larger size of PGCCs.

To verify the first possibility we measured the reactive oxygen species (ROS) production using H2DCF-DA fluorescent probe. Despite the progressively increased intracellular thiols and ROS-neutralizing enzymes, the antioxidant response was insufficient to fully counteract the accumulation of reactive oxygen species, which raised over time after doxorubicin treatment in parallel with the expanding PGCC population (Fig. 3E-F). This suggests that polyaneuploid transition and PGCC maturation are associated with elevated ROS production, which may activate antioxidant systems, and that observed increase in thiol content more likely results from the enlarged size of PGCCs. Possibly, PGCCs adapt to a distinct redox state in which elevated ROS levels are counterbalanced by increased thiol availability and some antioxidant enzymes, thereby maintaining conditions compatible with their survival.

In order to have an idea on the possible contribution of particular antioxidant factors identified by single-cell transcriptomics, we screened the impact of their inhibitors on PGCC survival (Fig. 3G) by making use of commercially available agents such as PX-12 to inhibit tioredoxin, Auranofin to Txnrd1 and, to lesser extent, Gpx1, NBDHEX to Gstp1 and Polaprezinc, which was recently reported to inhibit Prdx5 (38). Among the considered compounds, only inhibition of Txrd1/Gpx1 substantially declined viability of PGCC as measured by resazurin assay. A weaker, but not statistically significant effect was observed also for thioredoxin inhibitor. However, confocal imaging confirmed that both iTxn and iTxnrd1/Gpx1 inhibitors cause apoptotic and necrotic death of PGCC (Fig. 3H-I), as evidenced by annexin V and propidium iodide staining. Despite the more pronounced effect of iTxnrd1, both inhibitors substantially reduced intracellular thiol levels content in PGCC (Fig. 3J) and caused further increase in ROS level (Fig. 3K), the extent of which was comparable for both tested compounds.

In summary, the formation and maturation of PGCCs are associated with progressively increasing ROS production, which is accompanied by the induction of key antioxidant proteins functionally involved in maintaining redox homeostasis. PGCCs appear to tolerate sustained oxidative stress without compromising their viability, but pharmacological disruption of the thioredoxin system, which further shifts the intracellular redox balance toward a pro-oxidant state, results in the elimination of PGCCs.

Gauss distribution was tested with Shapiro-Wilk test. In (D), (F), (I), (J) and (K) sample variation was compared by ANOVA1 and posthoc Dunnett’s or Tukey’s test. In (G) statistical analysis was performed with Kruskal-Wallis and Dunn’s multiple comparison posthoc test. The difference was marked with * when adjusted p<0.05.

### 3.4 The Tnx–Txnrd1 system is essential for PGCC survival

Confocal imaging of tumor sections confirmed the elevated Txn and Txnrd1 levels in doxo-resistant MDA-MB-231 xenografts following six cycles of doxorubicin treatment, concomitant with an increased frequency of PGCCs (Fig. 4A-D). In contrast, tumors derived from cispl-resistant A549 (*TP53*^wt^) with relatively low frequency of PGCC after 6 cycles of chemotherapy with cisplatin, were characterized by statistically significant decline in expression of both proteins. The substantial increase in Txn and Txnrd1 was accordingly found in other *TP53*^mut^ drug-resistant cell lines exposed to doxorubicin for a period that allowed formation of PGCC, but not in *TP53^wt^* genotypes (Fig. 4E).

**Figure 4.**
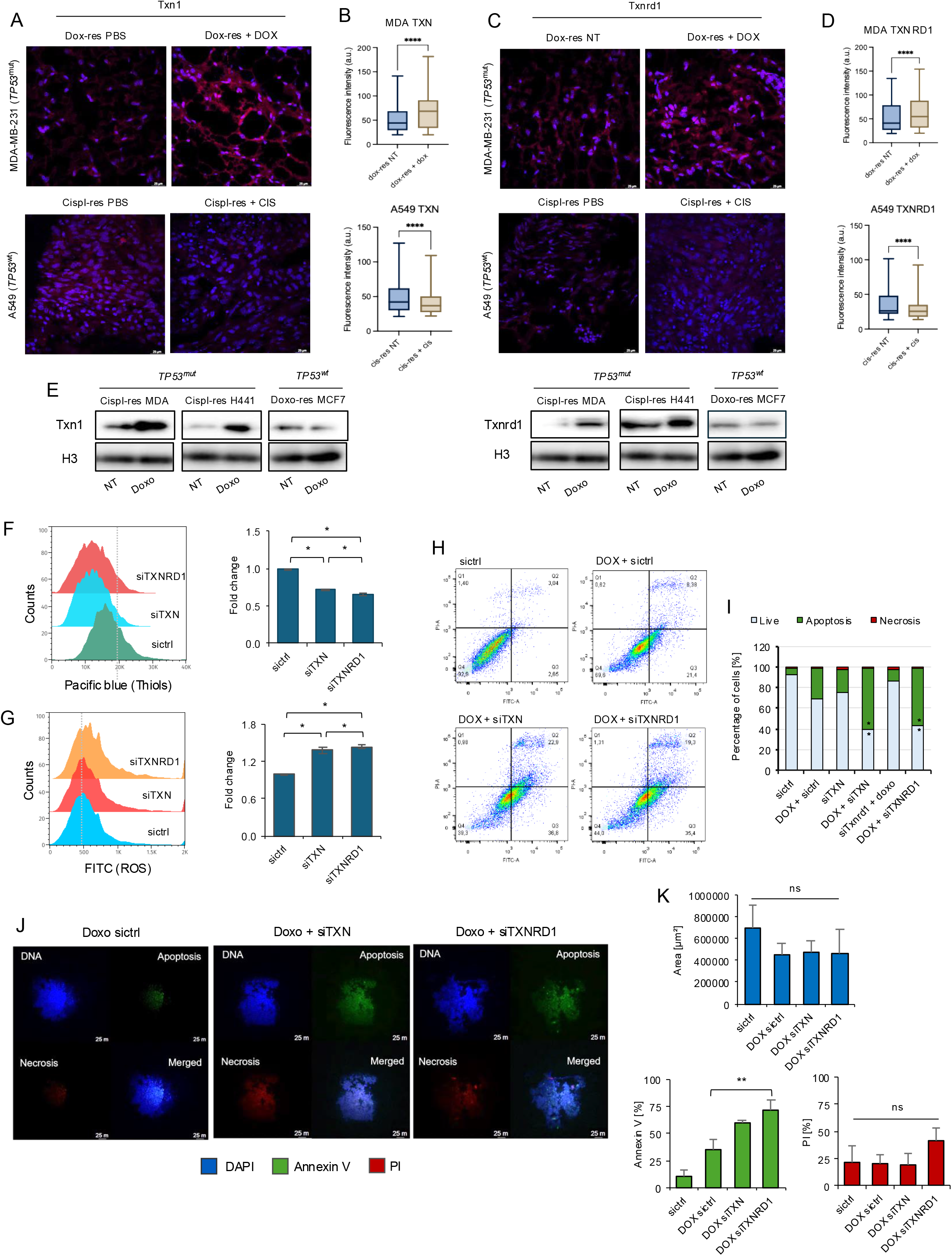
Thioredoxin – thioredoxin reductase 1 system protects PGCC from oxidative condition and preserves their viability (A-D) Confocal images confirm overexpression of Txn and Txnrd1 in *TP53^mut^* doxo-resistant, but not in *TP53^wild-type^* cispl-resistant tumors treated with 6 cycles of doxorubicin or cisplatin, respectively. In (A&C) thioredoxin and thioredoxin reductase 1 were stained with AlexaFluor546. (B and D) Their level was estimated by measuring fluorescence intensity with Leica Application Suite X - LAS X. (E) Western blot comparison of Txn and Txnrd1 level after doxorubicin treatment in *TP53^mut^* and *TP53^wt^* cell lines. (F-G) Thiol and ROS level was compared between PGCC with normal (sictrl), silenced *TXN* (si*TXN*) and *TXNRD1* (si*TXNRD*1) as in Fig. 3C-F. Cells were transfected with siRNA 24 h after doxorubicin, whereas their thiol (F) and ROS (G) level was measured after another 72 h. (H-I) The frequency of apoptotic and necrotic cells in the population of normal as well as *TXN* and *TXNRD1*-deficient PGCC was quantified by flow cytometry after double staining with annexin V (FITC) and propidium iodide (PE). Cell treatment scheme was the same as in (F-G). In (H) representative dotplots of double stained untreated cells and PGCC of selected phenotypes. (I) The average percentage of living, apoptotic and necrotic cells in the considered samples. (J-K) Apoptosis and necrosis in control and TXN- or TXNRD1-silenced spheroids were assessed using confocal microscopy using Annexin V–FITC and propidium iodide (PI), respectively. Fluorescence intensity was quantified using LAS X software. The treatment scheme was as same as in (F-G and H-I) (J) Representative images were acquired using a Leica LSI system. (K) The percentage of Annexin V- and PI-positive area quantified using LAS X software.

The transient silencing of *TXN* and *TXNRD1* with siRNA (Supplem. Fig. 3A) phenocopied the decline in thiol content, which was found in PGCC treated with iTxn or iTxnrd1/Gpx1 (Fig. 4F and Fig. 3J). Accordingly, the observed drop in thiols was followed by increased ROS, thereby indicating the shift to more oxidative condition (Fig. 4G). Notably, the deficiency of Txnrd1 resulted in a considerably more pronounced effect on thiols and ROS levels than silencing of *TXN*. Despite this marked difference, the silencing of *TXN* and *TXNRD1* induced similar extent of apoptosis in 2D culture of PGCC and reached approximately 40% (Fig. 4H,-4I and Supplem. Fig 3B-C), whereas the frequency of necrotic cells remained low. Some apoptosis was also observed in the culture of PGCC (sictrl + doxo; ∼20% vs sictrl), which are the mixture of early and late PGCC, but also cells that were not induced for polyaneuploid transition and die in response to doxorubicin.

To validate the effects of *TXN* and *TXNRD1* silencing in a more physiologically relevant tumor-like model, we employed three-dimensional spheroid cultures and transfected them with siTXN and siTXNRD1 24 h after their treatment with doxorubicin (Fig. 4J-K). The deficiency of Txn and Txnrd1 visibly increased the number of apoptotic cells, but quantification showed a significant increase only in siTXNRD1-transfected spheroids. Although the spheroid size remained unchanged, silencing of *TXN* and *TXNRD1* resulted in disrupted spheroid integrity, manifested by the appearance of fluorescence-free areas lacking labelled cell nuclei.

Depending on data distribution, differences between multiple groups were analysed using either a one-way ANOVA or a Kruskal–Wallis test and differences of two means was tested with unpaired t test with Welch’s correction. * means statistically significant when adjusted p < 0.05.

### 3.5 The Txn–Txnrd1 axis is associated with clinical outcome and anthracycline response in *TP53*^mut^ triple-negative breast cancer

Bearing in mind the critical role of the Txn–Txnrd1 system in PGCC and accumulating evidence from both the literature and our own studies that polyaneuploid transition occurs more often in *TP53*-mutant tumors, we expanded our analysis to include additional non-resistant and drug-resistant cell lines with either wild-type or mutant *TP53* (Supplem. Fig. 2B). Depletion of Txn or Txnrd1 was not directly cytotoxic, regardless of the genotype or phenotype of the cell lines examined. All three basal cell lines, which show only a limited capacity to undergo polyaneuploid transition, responded similarly to doxorubicin with substantial decline in their viability (Fig. 5A), but synergistic effect of *TXN* silencing and anthracycline was only observed in MDA-MB-231 cell line. On the contrary, the impact of *TXN* and *TXNRD1* silencing was genotype-dependent in drug-resistant cells, and an enhanced doxorubicin-induced cytotoxicity caused by si*TXN* or si*TXNRD1* was restricted to *TP53^mut^* cell lines.

**Figure 5.**
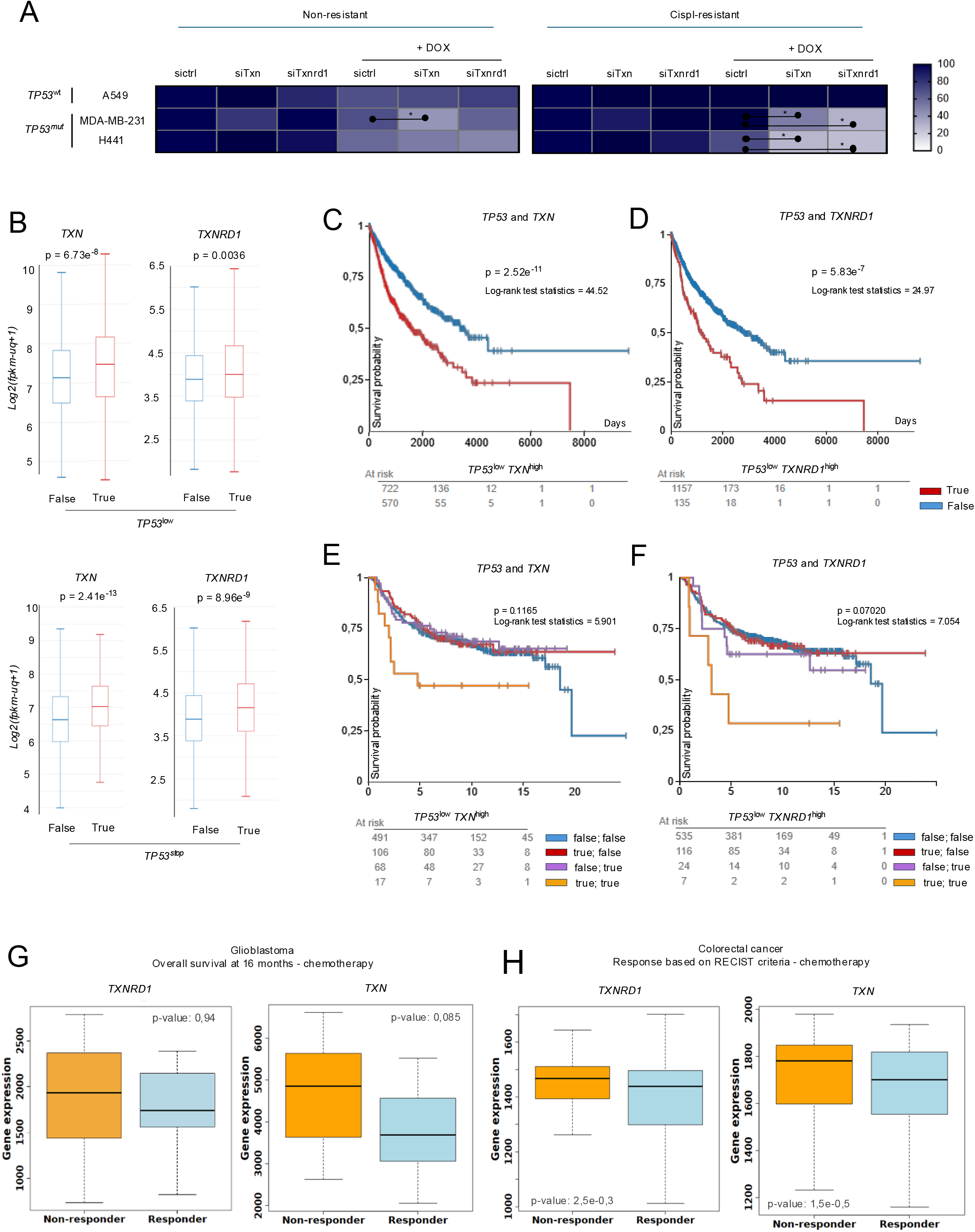
High *TXN* or *TXNRD1* expression in *TP53*-mutant or *TP53*-low TNBC may be associated with poor patient survival (A) The impact of Txn and Txnrd1 on viability of doxorubicin-treated cell lines with wild type and mutated *TP53*. Cell viability was measured by resazurin assay across cell genotypes and phenotypes. Cells were transfected 24 h after their treatment with doxorubicin. * indicates adj p < 0.05 according to Anova1 and Tukey post-hoc tests. (B) The impact of *TP53* expression or truncation (premature stop codon) in *TP53* on transcription of *TXN* and *TXNRD1* was tested in GDC Pan-Cancer clinical dataset. (C-D) Kaplan–Meier survival analysis of overall survival was performed using the same dataset for patients with low *TP53* expression in combination with high *TXN* (C) or *TXNRD1* (D). (E-F) Kaplan-Meier curves for distant metastasis free survival was created from Breast Cancer (Yau 2010) dataset for subgroups, which differed in expression level of *TP53* and *TXN* or *TXNRD1*. (G) The relationship between gene expression levels of *TXN* or *TXNRD1* and overall survival at 16 months in response to chemotherapy in patients with glioblastoma. (H) The association between *TXN* and *TXNRD1* gene expression levels and response to chemotherapy based on RECIST criteria in patients with colorectal cancer. The analyses (G-H) were performed using the ROC Plotter database.

In clinical datasets low expression of *TP53* or truncating mutation (premature stop codon) were linked to substantially higher transcription of *TXN* or *TXNRD1* (Fig. 5B). Probability of patient survival was significantly lower in patients with simultaneously high *TXN* or *TXNRD1* in combination with low *TP53* expression (Fig. 5C-D). Very similar tendency was observed for probability of distant metastases, where the Kaplan–Meier curves for tumors with concurrently low *TP53* and high *TXN* or T*XNRD1* expression diverged even more markedly from all other expression profiles (Fig. 5E-F). Notably, the combined *TP53*^low^–*TXNRD1*^high^ stratification showed the strongest prognostic trend (log-rank statistic = 7.054, p = 0.0702).

To evaluate the probably clinical relevance of *TXN* or *TXNRD1* in predicting response to chemotherapy, expression of both genes was compared between responders and non-responders (Fig. 5G-H). Non-responders with glioblastoma and colorectal cancer were characterized by significantly higher *TXN* and *TXNRD1* expression than responders (in glioblastoma p = 0,94 and p = 0,085, respectively and in colorectal cancer p = 2,5e-0,3 for TXNRD1 and p = 1,5e-0,5 for TXN). Unfortunately, due to methodological limitations in such analyses, *TP53* expression levels and mutational status could not be incorporated.

In summary, these findings are consistent with our experimental observations, suggesting that elevated Txn–Txnrd1 activity may be particularly detrimental in *TP53*-low or -deficient tumors, where PGCC formation is more prevalent and may contribute to disease progression and poor clinical outcome.

## Discussion

Our single-cell RNA sequencing results uncovered the heterogeneity of PGCCs and allowed us to characterize transcriptomes of early and mature PGCC. In addition to metabolic reprogramming, which made these cells more dependent on oxidative phosphorylation as also described in former experimental papers, we provided evidence on adaptive redox remodelling. By monitoring cells throughout polyaneuploid transition and subsequent PGCC maturation, we found that the progressive increase in reactive oxygen species was accompanied by a coordinated upregulation of antioxidant defence mechanisms. This adaptive response involved increased expression of thioredoxin, together with enzymes that rely on the thioredoxin and glutathione systems to detoxify reactive oxygen species and restore oxidized protein thiols. Similar phenomenon was described in other cancer models and was assigned to the mechanism of redox non-oncogene addiction, where elevated reactive oxygen species force reliance on antioxidant buffering systems to prevent cell death (39–41). Conceptually, our model can be described by the following sequence of temporal changes: doxorubicin -> increased ROS levels -> polyaneuploid transition -> adaptive upregulation of *TXN/TXNRD1* and other antioxidant systems -> survival under persistent oxidative stress -> acquired redox dependency/non-oncogene addiction, where the stress-adapted phenotype renders cancer cells disproportionately dependent on the thioredoxin system for survival. Such a dependency can be therapeutically exploited to enhance the efficacy of chemotherapy, as demonstrated by our experiments involving genetic silencing and pharmacological inhibition of Txn or Txnrd1, as well as by previous studies conducted in other *in vitro* and preclinical cancer models (39,42,43).

Although doxorubicin is known to increase intracellular ROS levels (44), the first detectable shift in redox balance toward a more oxidative state coincided with PGCC formation in our model (Fig. 4E–F), whereas a marked increase in cellular thiol content was evident 24 h after doxorubicin treatment. This temporal discrepancy may reflect an early compensatory antioxidant response, in which the intracellular thiols pool buffers doxorubicin-induced ROS and delays a measurable shift toward a more oxidative redox state, and progressively increased oxidative pressure leads to adaptive redox rewiring with thiol-dependent antioxidant enzymes overexpression. Earlier studies have implicated mitochondria as a major source of ROS in PGCCs (18,45), and some linked high ROS levels to increased tumorigenicity and resistance to paclitaxel. However, the reported increment of lactic acid as well as activation of pentose phosphate pathway in PGCCs may suggest that these cells take the advantage of both aerobic and anaerobic pathways for the energy production, but such hypothesis requires verification in future studies (17,45). Gene ontology analysis of genes upregulated in early and mature PGCCs revealed enrichment of mitochondrial ATP production and oxidative phosphorylation, particularly associated with complex 1 and 3 in mitochondrial respiratory chain, thereby suggesting that metabolic rewiring is initiated early during the polyaneuploid transition and persists throughout PGCC maturation. Nevertheless, these findings do not exclude the involvement of additional metabolic pathways that may contribute to the metabolic plasticity and survival of PGCC. Since our single-cell transcriptomic data did not reveal increased expression of NOX1–5, DUOX1/2, xanthine oxidoreductase, peroxisomal oxidases, or cytochrome P450 enzymes, the mitochondrial electron transport chain emerges as the most plausible source of the increased ROS observed in PGCC. But also here, further experimental studies are needed.

Such a strong and direct dependence of PGCCs on the Txn–Txnrd1 system may appear unexpected, particularly given that Txn is not the predominant intracellular antioxidant. Earlier studies demonstrated compensatory mechanisms between the glutathione and thioredoxin systems and, in some contexts, even synthetic lethality, where cells with low glutathione reductase expression were particularly vulnerable to inhibition of the Txn–Txnrd1 axis (46). More recent findings suggest that these two antioxidant networks are not fully redundant; rather, they exhibit distinct substrate specificities and regulate partially non-overlapping sets of redox-sensitive proteins and signalling pathways (47,48). In addition to its antioxidant function, the Txn-Txnrd1 system plays a much broader range of biological roles being involved in the regeneration of peroxiredoxins, synthesis of DNA by supplying electrons to ribonucleotide reductase, maintaining conserved Cys residues of transcription factors (NF-κB, AP-1, p53, Nrf2, glucocorticoid receptor (GR), estrogen receptor (ER), and HIF1) in the reduced form required for DNA binding by reducing redox factor-1 (Ref-1), and controlling pathway for MSRB2 reduction mediated by metallothionein (49). Prasad et. al. demonstrated that Txn/Txnrd1 maintains ribonucleotide reductase activity through redox recycling of its RRM1 subunit and, hence, sustains the dNTP pool required for DNA replication (50). Disruption of this system resulted in RRM1 oxidation, dNTP depletion, impaired replication fork progression and consequent replication stress and DNA damage. This mechanism may be particularly relevant to PGCCs, which undergo extensive DNA synthesis during the polyaneuploid transition and presumably impose an unusually high demand on nucleotide biosynthesis. In the context of the potential contribution of GSH to PGCC function under shifter redox balance, our single-cell data did not indicate increased expression of enzymes involved in glutathione biosynthesis or glutathione reductase. Instead, the transcriptional profile pointed toward a preferential role of GSH in GST-mediated drug conjugation, as two glutathione S-transferases were upregulated in PGCCs. Consistent with this interpretation, GSTP1 inhibition did not compromise PGCC viability, likely because the drug was no longer present in the extracellular environment at the later stages of PGCC formation. Moreover, the pre-existing glutathione pool in PGCCs may have been sufficient to sustain the newly established redox homeostasis without requiring increased GSH biosynthesis or recycling.

Despite the absence of detectable *TXN2* and *TXNRD1* mRNA upregulation, the protein levels of both enzymes were increased in PGCCs. This suggests that the abundance of these proteins is regulated at the post-transcriptional level and may result from enhanced translation or increased protein stability due to reduced proteasomal degradation under stress conditions (51,52). Mitochondrial Txn2 primarily supports local mitochondrial redox homeostasis and may play a less prominent role in PGCC survival than the cytosolic Txn1–Txnrd1 system, which operates in cytoplasm and nucleus (49). Previous studies have extensively implicated Txnrd1 in cancer cell physiology and identified this enzyme as a potential therapeutic target, inhibition of which was expected to provide a degree of selectivity toward malignant cells (53). Interestingly, in some cases, Txnrd1 inhibition converted the enzyme into pro-oxidant SecTRAPs with NADPH oxidase activity, thereby exacerbating oxidative stress, particularly in cells with high Txnrd1 expression (54). According to Patwardhan et al. Txnrd1 was not over-expressed at mRNA or protein level in breast primary tumor but was considerably overexpressed in hormone receptor-negative and HER2-positive tumors and showed positive correlation with tumor grade and size (55). Moreover, Txnrd1 over-expression indicated a poor prognosis in breast cancer patients, but the Authors did not account for *TXN* expression, precluding assessment of the prognostic relevance of the complete Txn1–Txnrd1 axis. The observation on *TXNRD1* alone is consistent with our, who were further stratified according to *TP53* expression, with low *TP53* levels representing a molecular context associated with increased predisposition to polyaneuploid transition. Our analyses suggest that the adverse prognostic impact of high *TXN1* or *TXNRD1* expression on patient survival and distant metastasis becomes apparent specifically in the context of low *TP53* expression. Furthermore, across the drug-resistant non-small cell lung cancer models, drug-induced upregulation of *TXN1* and *TXNRD1* was associated with *TP53* status and with the ability of cancer cells to undergo polyaneuploid transition, rather than with ER, PGR, or HER2 receptor expression. Although the polyaneuploid transition is not determined by *TP53* status alone but appears to be strongly dependent on the acquisition of chemoresistance, we found the strong association between expression of *TP53* and *TXN1* or *TXNRD1* in clinical datasets. Unfortunately, further stratification according to acquired therapy resistance cannot be reliably performed using currently available large-scale clinical datasets, which could possibly reveal even stronger interdependence between *TP53* and *TXN1* or *TXNRD1.* Importantly, higher expression of both genes was also associated with chemoresistance in the intestinal cancer and glioblastoma. At least some of the latter tumors with mutation in *TP53* were documented to form PGCC after their exposure to temozolomide and PARP inhibitor (34,35).

Concluding, Txn1-Txnrd1 system confers PGCC viability under rewired redox homeostasis and pharmacological targeting of Txn1 or Txnrd1 may be considered as new strategy candidate to overcome chemoresistance in cancers with low or damaging mutations in *TP53*.

## Supporting information

Supplementary Results Figures

Supplementary Results Legends

Supplementary Results Western Blot

## Acknowledgments

IDUB Doctoral Research Grant 2024 edition; STSM COST CA17140 action; BioMedChem Doctoral School of University of Lodz and Institutes of Polish Academy of Sciences

---

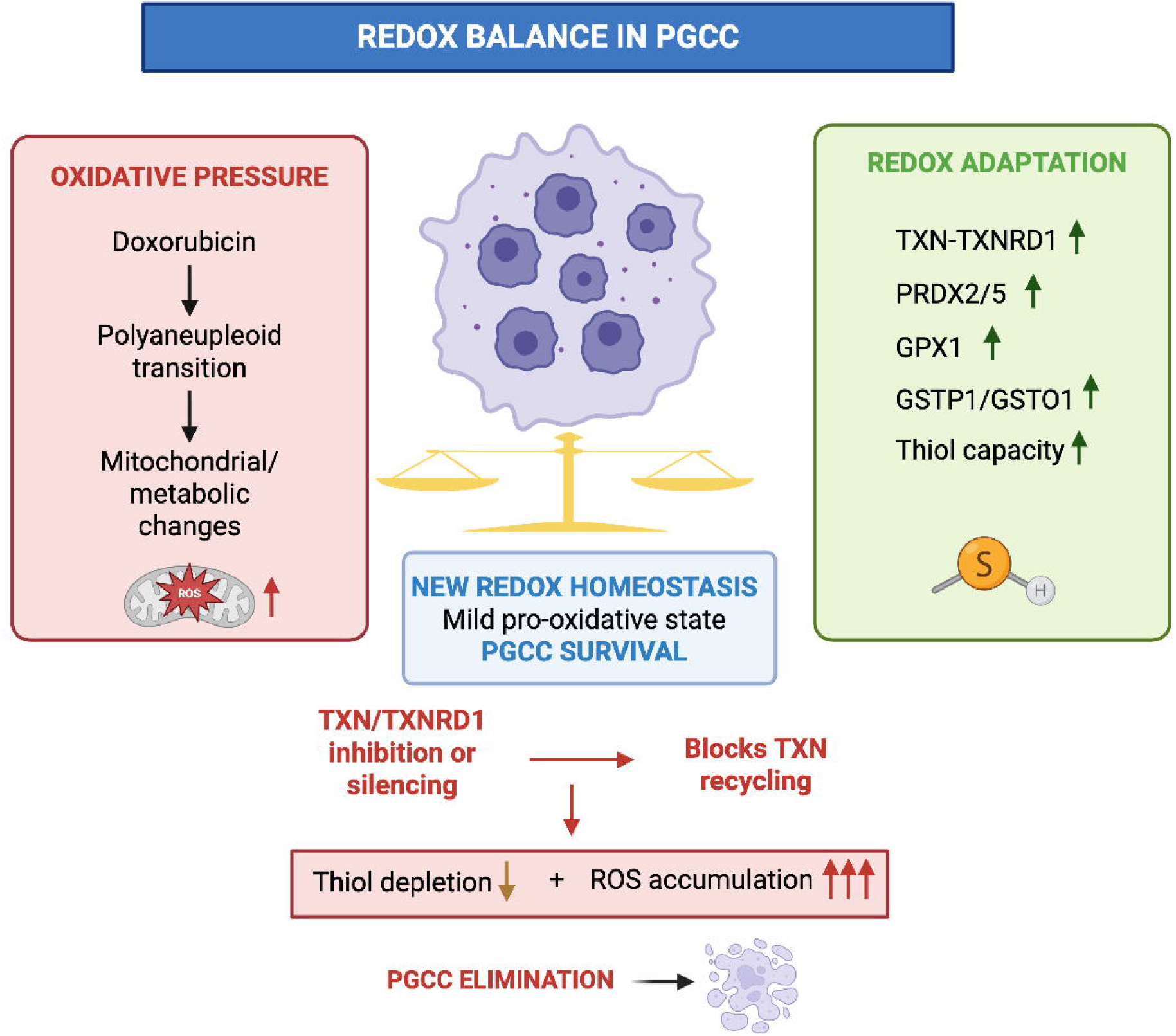

## Notes

### Competing Interest Statement

The authors have declared no competing interest.

### Summary of Updates

Figures 3 and 5 have been updated to correct a minor labeling/typographical error.

