## Supplementary Results Figures for "Txn-Txnrd1 system supports redox rewiring during polyaneuploid transition and protects giant cancer cells at new redox homeostasis"

### Slide 1
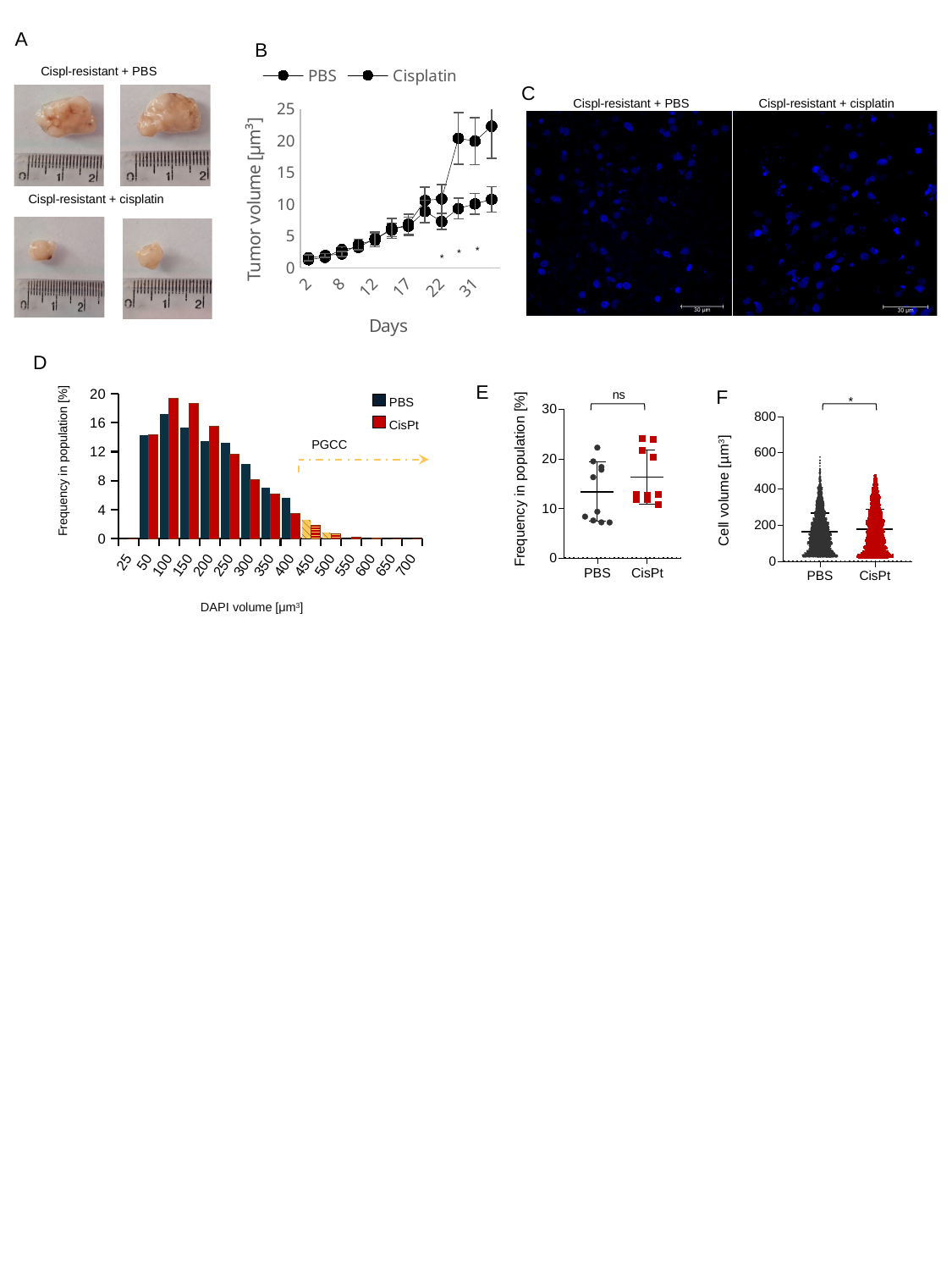

A
B
Cispl-resistant + PBS
Cispl-resistant + cisplatin
#### Chart
| Category | | |
|---|---|---|
| 2 | 1.266762440225431 | 1.55347205559624 |
| 5 | 1.6488017578336804 | 1.91554845850125 |
| 8 | 2.854514674289255 | 2.1859708520530807 |
| 10 | 3.2376224144032206 | 3.68458362031918 |
| 12 | 4.607489368675056 | 4.385127378111082 |
| 15 | 5.924885510826319 | 6.200243834898233 |
| 17 | 6.866964828269312 | 6.542073829032096 |
| 19 | 10.601462449643892 | 8.875834101439226 |
| 22 | 10.860918847141361 | 7.2808982035043135 |
| 29 | 20.4077240715122 | 9.336846972501831 |
| 31 | 19.949844468431845 | 10.06404329648647 |
| 36 | 22.289986183353935 | 10.781925223416277 |*
*
*
C
Cispl-resistant + PBS
Cispl-resistant + cisplatin
D
#### Chart
| Category | | |
|---|---|---|
| 25 | 0.0 | 0.0 |
| 50 | 14.323911382734913 | 14.29685976204408 |
| 100 | 17.265087853323145 | 19.348546908523502 |
| 150 | 15.355233002291827 | 18.704895650477862 |
| 200 | 13.445378151260504 | 15.54515311098108 |
| 250 | 13.254392666157372 | 11.683245562707237 |
| 300 | 10.313216195569137 | 8.152915935244783 |
| 350 | 7.066462948815889 | 6.104934659645016 |
| 400 | 5.576776165011459 | 3.3937975424224693 |
| 450 | 2.559205500381971 | 1.8724400234055003 |
| 500 | 0.8403361344537815 | 0.6436512580456407 |
| 550 | 0.0 | 0.19504583577140627 |
| 600 | 0.0 | 0.058513750731421885 |
| 650 | 0.0 | 0.0 |
| 700 | 0.0 | 0.0 |ns
Frequency in population [%]
E
F
*
PBS
CisPt
Cell volume [µm^3^]
PGCC
Frequency in population [%]
DAPI volume [μm^3^]

### Slide 2
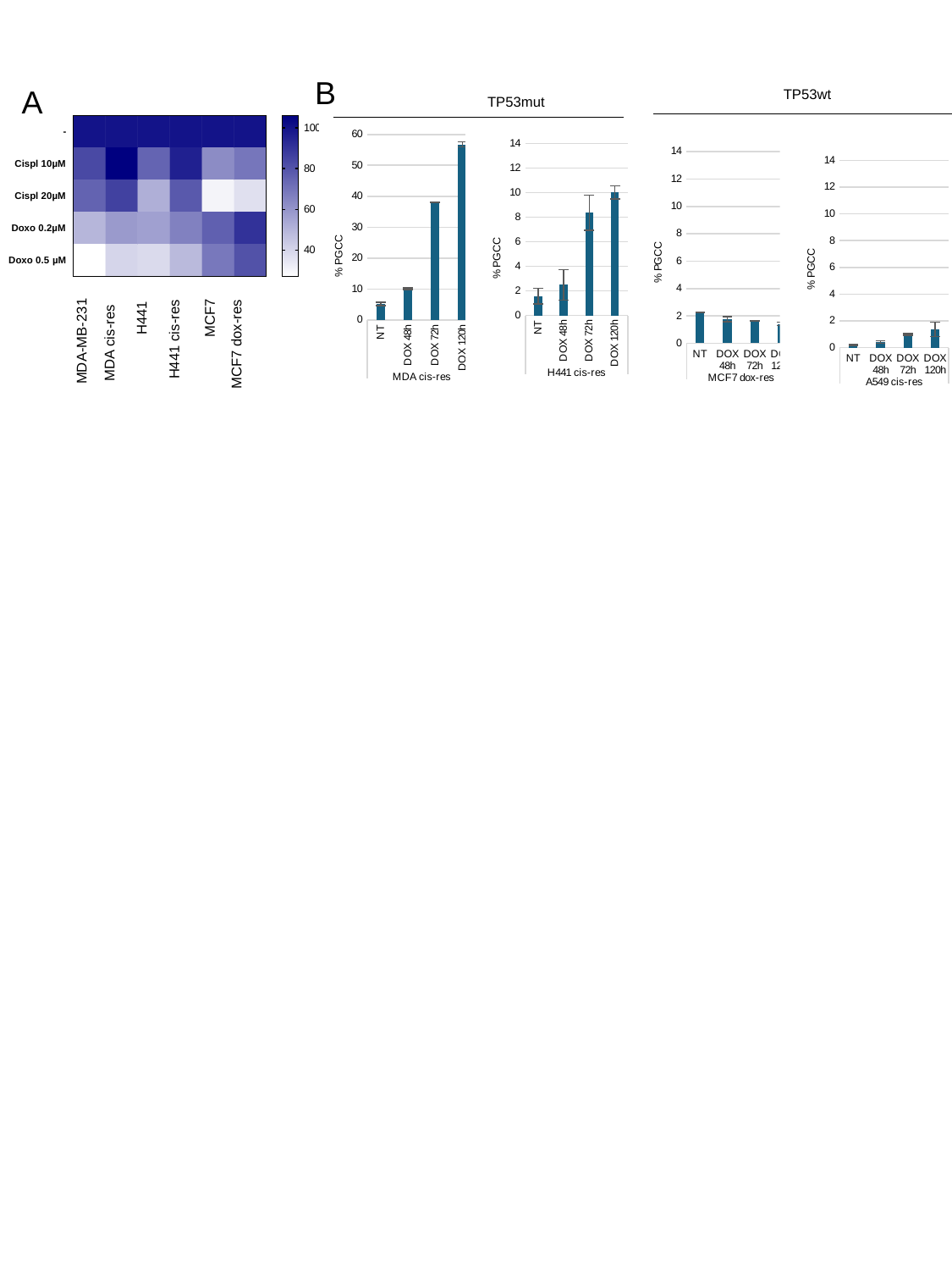

B
A
TP53wt
TP53mut
#### Chart
| Category | |
|---|---|
| NT | 5.125 |
| DOX 48h | 10.105 |
| DOX 72h | 37.95 |
| DOX 120h | 56.75 |
#### Chart
| Category | |
|---|---|
| NT | 1.585 |
| DOX 48h | 2.49 |
| DOX 72h | 8.365 |
| DOX 120h | 10.025 |
#### Chart
| Category | |
|---|---|
| NT | 2.265 |
| DOX 48h | 1.75 |
| DOX 72h | 1.6549999999999998 |
| DOX 120h | 1.4100000000000001 |
#### Chart
| Category | |
|---|---|
| NT | 0.175 |
| DOX 48h | 0.42 |
| DOX 72h | 0.985 |
| DOX 120h | 1.365 |H441
MCF7
H441 cis-res
MDA cis-res
MDA-MB-231
MCF7 dox-res

### Slide 3
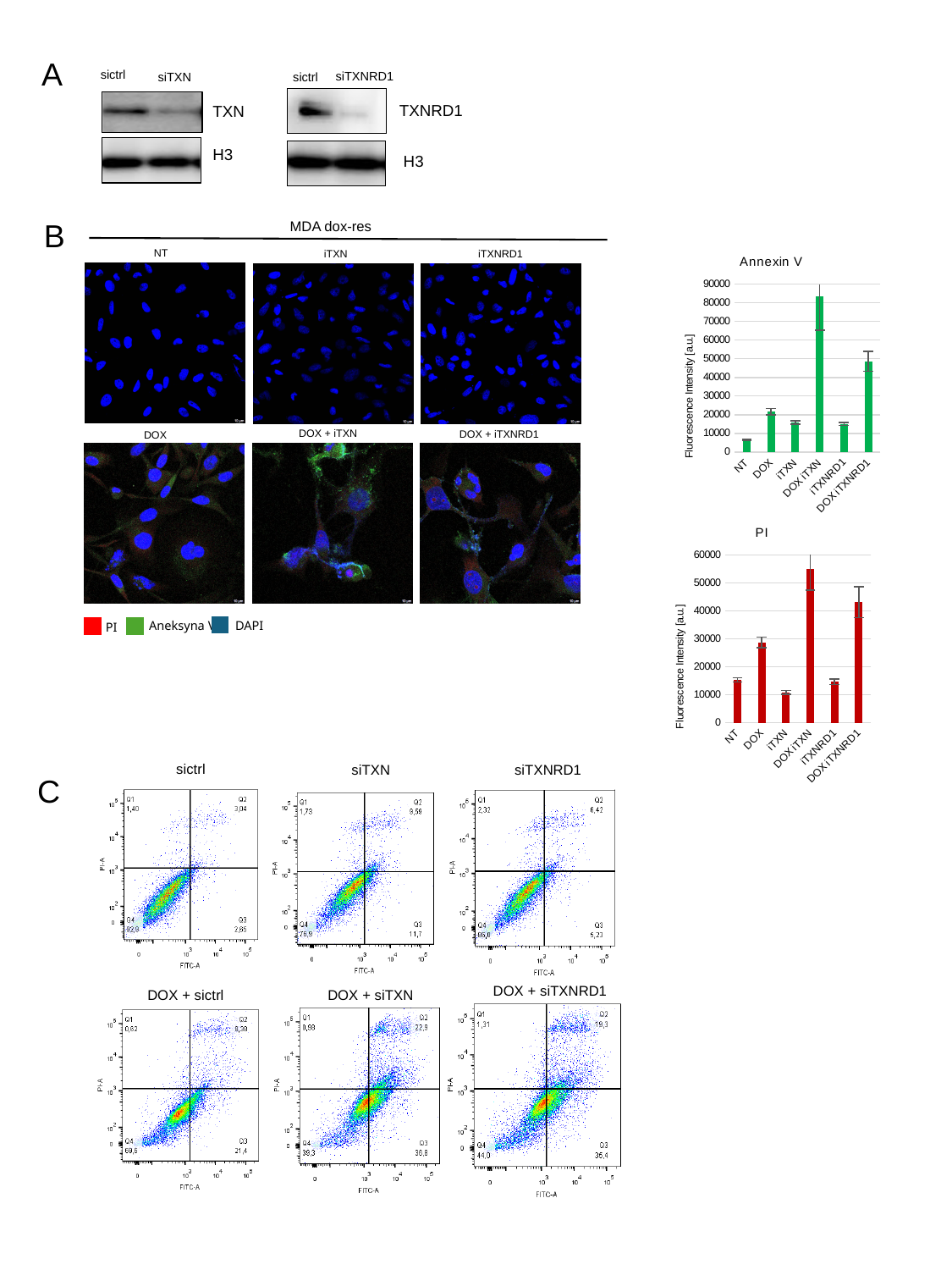

A
sictrl
siTXNRD1
siTXN
sictrl
TXNRD1
TXN
H3
H3
B
MDA dox-res
#### Chart: Annexin V
| Category | |
|---|---|
| NT | 6467.9 |
| DOX | 21582.61111111111 |
| iTXN | 15877.08695652174 |
| DOX iTXN | 83241.86666666667 |
| iTXNRD1 | 14939.058536585366 |
| DOX iTXNRD1 | 48594.333333333336 |NT
iTXNRD1
iTXN
DOX + iTXN
DOX + iTXNRD1
DOX
#### Chart: PI
| Category | |
|---|---|
| NT | 15323.05 |
| DOX | 28676.055555555555 |
| iTXN | 10760.652173913044 |
| DOX iTXN | 55044.26666666667 |
| iTXNRD1 | 14671.388780487809 |
| DOX iTXNRD1 | 43083.333333333336 |DAPI
Aneksyna V
PI
sictrl
siTXN
siTXNRD1
C
DOX + siTXNRD1
DOX + sictrl
DOX + siTXN
