## Supplementary Results Legends for "Txn-Txnrd1 system supports redox rewiring during polyaneuploid transition and protects giant cancer cells at new redox homeostasis"

Supplementary Fig 1. Negligible formation of giant polyaneuploid cells (PGCC) in xenografts formed from A549 cells

(A)Representative images of tumors grown and isolated from athymic mice treated with PBS or cisplatin (10 mg/kg body weight). (B) Tumor growth kinetics (mm³) following PBS or cisplatin treatment administered every 7 days. (C) Representative fluorescence images of tumor sections from PBS- or drug-treated mice. Nuclei are stained with DAPI. (D) Distribution of DAPI-stained nuclear volumes determined from confocal microscopy images. Cells with nuclear volume > 3xG1 (highest peak in PBS injected mice) were assumed as PGCC and marked in orange. Quantification of PGCC frequency (E) and nuclear volume (F) in tumor sections.

Suplementary Fig 2. PGCC formation in populations of chemotherapy-resistant cells with mutant and wild-type p53

(A)The acquisition of drug resistance in MDA-MB-231, NCI-H441 and MCF7 cell lines was determined by comparing their viability in response to two concentrations of selected drugs, which were added to cell culture for 48 h. Drug toxicity was quantified by resazurin assay, and the fluorescence of untreated cells was assumed as 100%.(B) The percentage of PGCCs in populations of chemotherapy-resistant cells with mutant (MDA-MB-231 and H441) or wild-type (A549 and MCF7) p53 following exposure to doxorubicin. The percentage of PGCCs was determined by analysing the DNA content of MDA-MB-231, H441, MCF7 and A549 cells using flow cytometry, under control conditions (NT) and after 48, 72 and 120 hours of exposure to doxorubicin (DOX). The cells were resistant to either cisplatin or doxorubicin, as indicated. Data are presented as mean ± SD.

Supplementary Fig 3 The Tnx–Txnrd1 system is essential for PGCC survival

(A) Western blot analysis of TXN and TXNRD1 in the MDA-MB-231 cell line, in the NT state and following TXN or TXNRD1 silencing. (B) Representative confocal images of MDA-MB-321 cells before and after doxorubicin treatment, as well as with TXN or TXNRD1 inhibition alone or in combination with doxorubicin. Nuclei are stained with DAPI and apoptotic cells with annexin V-FITC, while necrotic cells are stained with propidium iodide (PI). The charts on the right show the quantitative analysis of fluorescence intensity expressed in arbitrary units. (C) Flow cytometry plots showing apoptosis and necrosis in MDA-MB-231 cells following the indicated treatments. The cells were double-stained with Annexin V-FITC and propidium iodide (PI). The quadrants represent the following: viable cells (Annexin V⁻/PI⁻; Q4); early apoptotic cells (Annexin V⁺/PI⁻; Q3); late apoptotic cells (Annexin V⁺/PI⁺; Q2); and necrotic cells (Annexin V⁻/PI⁺; Q1). The numbers indicate the percentage of cells in each group.
