## Supplementary Results Western Blot for "Txn-Txnrd1 system supports redox rewiring during polyaneuploid transition and protects giant cancer cells at new redox homeostasis"

### Slide 1
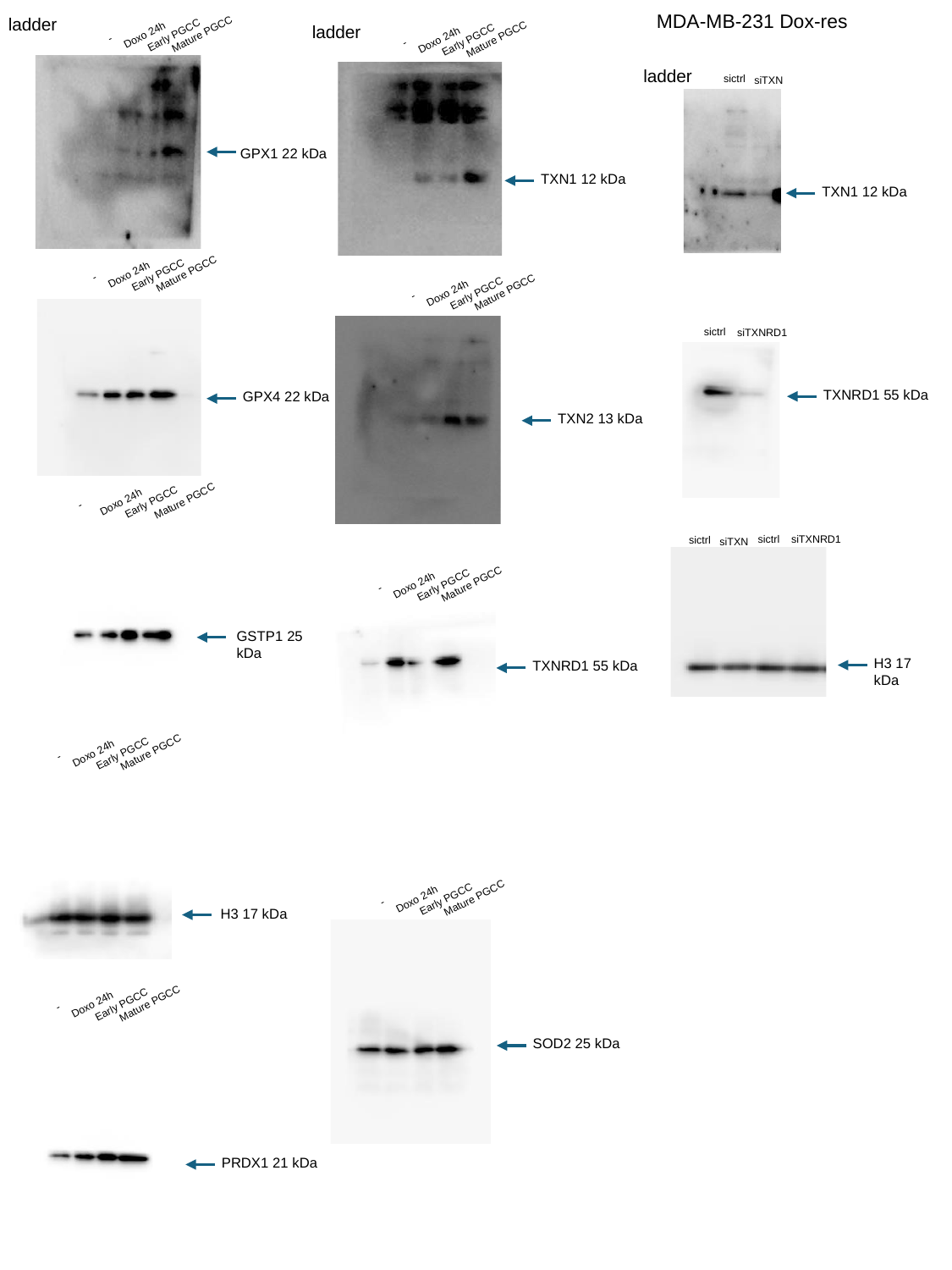

MDA-MB-231 Dox-res
ladder
ladder
Mature PGCC
Doxo 24h
Early PGCC
-
Mature PGCC
Doxo 24h
Early PGCC
-
ladder
sictrl
siTXN
GPX1 22 kDa
TXN1 12 kDa
TXN1 12 kDa
Mature PGCC
Doxo 24h
Early PGCC
-
Mature PGCC
Doxo 24h
Early PGCC
-
sictrl
siTXNRD1
TXNRD1 55 kDa
GPX4 22 kDa
TXN2 13 kDa
Mature PGCC
Doxo 24h
Early PGCC
-
sictrl
siTXNRD1
sictrl
siTXN
Mature PGCC
Doxo 24h
Early PGCC
-
GSTP1 25 kDa
H3 17 kDa
TXNRD1 55 kDa
Mature PGCC
Doxo 24h
Early PGCC
-
Mature PGCC
Doxo 24h
Early PGCC
-
H3 17 kDa
Mature PGCC
Doxo 24h
Early PGCC
-
SOD2 25 kDa
PRDX1 21 kDa

### Slide 2
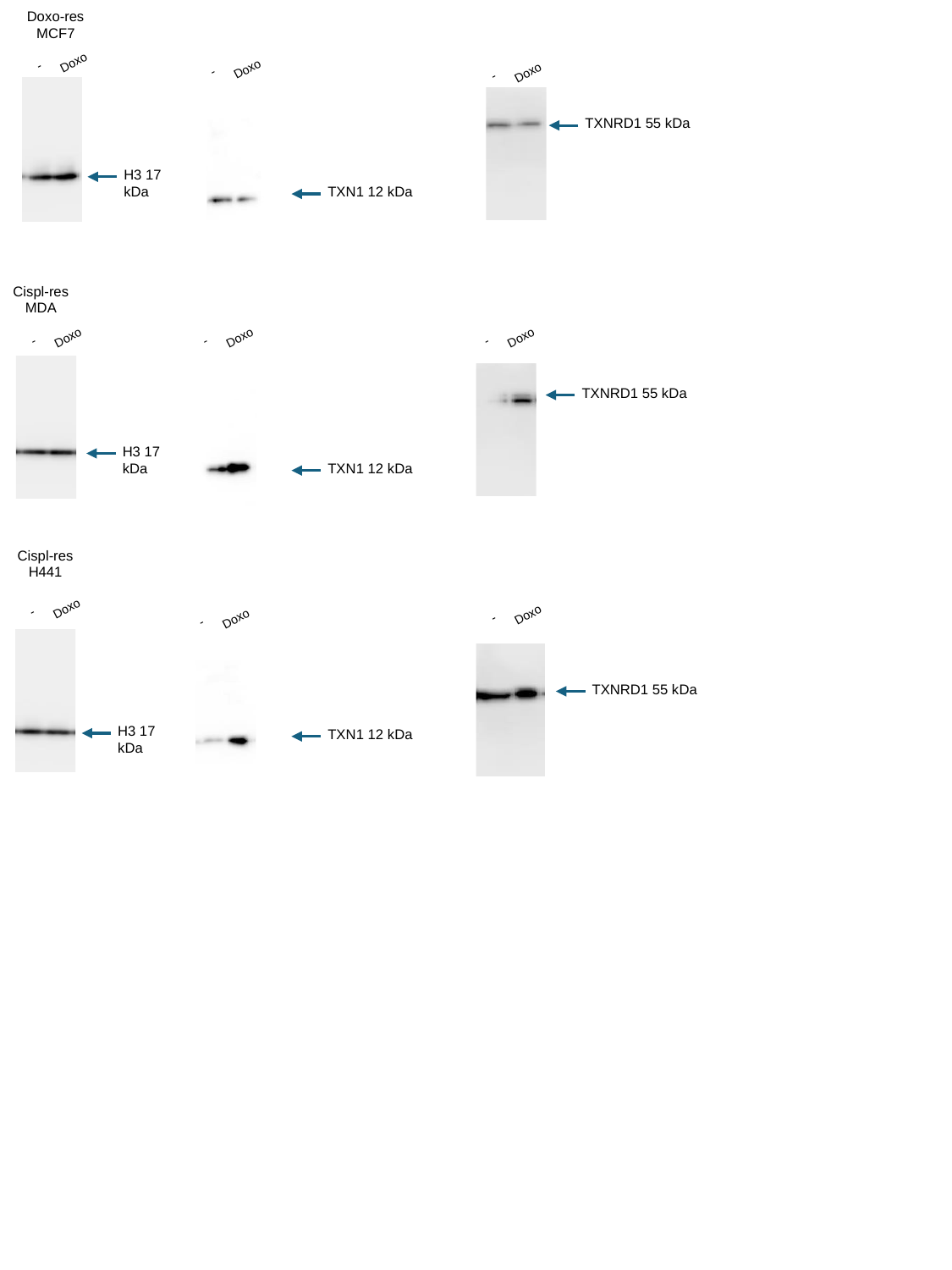

Doxo-res MCF7
Doxo
-
Doxo
-
Doxo
-
TXNRD1 55 kDa
H3 17 kDa
TXN1 12 kDa
Cispl-res MDA
Doxo
Doxo
Doxo
-
-
-
TXNRD1 55 kDa
H3 17 kDa
TXN1 12 kDa
Cispl-res H441
Doxo
-
Doxo
-
Doxo
-
TXNRD1 55 kDa
H3 17 kDa
TXN1 12 kDa
